# Probabilistic modeling methods for cell-free DNA methylation based cancer classification

**DOI:** 10.1101/2021.06.18.444402

**Authors:** Viivi Halla-aho, Harri Lähdesmäki

## Abstract

**Background:** cfMeDIP-seq is a low-cost method for determining the DNA methylation status of cell-free DNA and it has been successfully combined with statistical methods for accurate cancer diagnostics. We investigate the diagnostic classification aspect by applying statistical tests and dimension reduction techniques for feature selection and probabilistic modeling for the cancer type classification, and we also study the effect of sequencing depth.

**Methods:** We experiment with a variety of statistical methods that use different feature selection and feature extraction methods as well as probabilistic classifiers for diagnostic decision making. We test the (moderated) t-tests and the Fisher’s exact test for feature selection, principal component analysis (PCA) as well as iterative supervised PCA (ISPCA) for feature generation, and GLMnet and logistic regression methods with sparsity promoting priors for classification. Probabilistic programming language Stan is used to implement Bayesian inference for the probabilistic models.

**Results and conclusions:** We compare overlaps of differentially methylated genomic regions as chosen by different feature selection methods, and evaluate probabilistic classifiers by evaluating the area under the receiver operating characteristic (AUROC) scores on discovery and validation cohorts. While we observe that many methods perform equally well as, and occasionally considerably better than, GLMnet that was originally proposed for cfMeDIP-seq based cancer classification, we also observed that performance of different methods vary across sequencing depths, cancer types and study cohorts. Overall, methods that seem robust and promising include Fisher’s exact test and ISPCA for feature selection as well as a simple logistic regression model with the number of hyper and hypomethylated regions as features.

## Background

In recent years the interest in utilizing circulating cell-free DNA (cfDNA) for cancer diagnostics has grown, enhanced by the development of next-generation sequencing (NGS) technologies. Cell-free DNA refers to DNA fragments that are not associated with cells and is considered to originate from cell apoptosis and necrosis [1, 2]. In the case of a presence of a tumor, it can be the source of part of the cfDNA. cfDNA released by a tumor is called circulating tumor DNA (ctDNA). Cell-free DNA can be extracted in a minimally non-invasive manner from a bodily fluid sample, such as blood, to identify and detect cancer-type specific biomarkers [3].

ctDNA is believed to represent the tumor burden and to carry the same genomic and epigenetic properties as the tumor of origin [3]. Therefore multiple types of cancer biomarkers can be identified and detected from cfDNA, including mutations, epigenetic modifications and copy-number alterations [3]. The DNA fragmentation profiles of cfDNA can also be used to classify cancer types [4]. Compared to quantification of somatic mutations from sequencing data which necessarily requires a high sequencing coverage, using methylation biomarkers can significantly reduce costs. Consequently, in this work we concentrate on cancer classification which is based on DNA methylation biomarkers. The most common way to measure DNA methylome is bisulfite sequencing (BS-seq), and tools such as CancerLocator [5] utilize BS-seq data to learn machine learning models to classify different cancer types. However, bisulfite conversion step of the BS-seq method leads to high degradation of the input DNA [6], making it unsuitable for cfDNA analysis where the amount of sequencing material is small.

Cell-free methylated DNA immunoprecipitation and high-throughput sequencing (cfMeDIP-seq) is a protocol for measuring the methylation status of cell-free DNA [7]. cfMeDIP-seq is a version of the MeDIP-seq method that takes into account specific needs of cfDNA sequencing. The amount of cfDNA material available for sequencing is often very small, so filler DNA from *Enterobacteria phage λ* is used in cfMeDIP-seq to increase the amount of DNA material [7]. Compared to bisulfite sequencing, cfMeDIP-seq is even more cost-effective, as only the methylated reads are sequenced in the immunoprecipitation-based approach [8]. While bisulfite sequencing provides information of the methylation status in a single-base resolution, cfMeDIP-seq can give information on the methylation status of genomic regions of length around 100bp or more [8].

Along with the cfMeDIP-seq protocol, statistical methods for finding differentially methylated regions (DMRs) and machine learning methods for classification of the cancer types were proposed in [7]. In brief, DMRs are found for each cancer type and healthy controls (i.e., a class) using a moderated t-test that separately compares each class to other classes in one-vs.one manner. Then binary GLMnet [9] classifiers are trained for each of the classes, using the DMRs found in the previous step. The results of these methods presented in [7] show high accuracy both in the discovery and validation data cohorts. The results for renal cell carcinoma (RCC) class were further validated in [10], where the same methods were applied to classify RCC patients from healthy controls. The cfMeDIP-seq assays and analysis steps were performed not only for plasma cfDNA, but also for cfDNA of urine origin. Both resulted in high AUROC scores, although the plasma-based classifier performed slightly better. In [11], the cfMeDIP-seq data set from [7] was extended with samples from intracranial tumor patients.

There are also other works reporting usage of cfMeDIP-seq or MeDIP-seq measurements of cfDNA for cancer classification [12, 13, 14, 15]. Similar to [7], GLMnet models were utilized to classify pancreatic cancer patients and healthy controls in [12], but the model features were based on both cfMeDIP-seq and cell-free 5hmC sequencing data. Peak calling was performed for both cfMeDIP-seq and cell-free 5hmC signals with MACS2 tool [16], and differential peaks between the cancer samples and healthy controls were then determined with t-test. Both 5mC and 5hmC peaks used separately as model features gave high prediction accuracies, but using both peak types together yielded even better results. In [13], the performances of detecting metastatic renal cell carcinoma (mRCC) using cfMeDIP-seq based cfDNA methylation analysis and cfDNA variant analysis were compared. The TMM-normalized cfMeDIP-seq count data was used to find DMRs with limma-voom [17] and the DMRs were then utilised as features in a GLMnet model, similar to the approach in [7]. The comparison showed that the classification method based on cfMeDIP-seq data had considerably higher sensitivity than cfDNA variant analysis. MeDIP-seq has also been applied to small number of cfDNA samples to find DMRs between cancer patients and healthy individuals, in particular for lung cancer [14] and pancreatic cancer [15]. In these cases the DMRs were found using MEDIPS tool [18] and they were further used for analyzing methylation data of tissue origin. MEDIPS is a tool for quality control and analysis of immunoprecipitation sequencing data, and it performs differential coverage analysis using negative binomial model from edgeR package [19].

The results in [7] showed that methylation-based cfDNA biomarkers have a great potential in cancer classification and that cfMeDIP-seq is a sensitive yet low-cost method for measuring the methylome. However, if the cfMeDIP-seq method was to be applied in clinical use, we hypothesize that for enhancing cost-efficiency the sequencing depth would have to be lower than in the demonstrative data set shown in [7]. But how well would the classification methods presented in [7] cope with lower sequencing depth? In this work we attempt to simulate a situation where the sequencing depth is considerably lower. Additionally, we present statistical methods for improving the feature selection and probabilistic modeling to improve the classification of the cancer types. We compare our approaches to the machine learning methods presented in [7]. For feature selection, we experiment with classical principal component analysis (PCA) and iterative supervised PCA (ISPCA) [20], which can utilize the class information for finding the optimal principal components for separating classes from each other. We also apply Fisher’s exact test for DMR finding, as a simpler statistical test could be more robust when the sequencing depth is lower. For the classification methods, we experiment with logistic regression with regularized horseshoe (RHS) prior [21] and logistic regression with DMR count variables, both implemented with probabilistic programming language Stan [22].

## Methods

### Aim of the study

The aim of this study was to design and test various feature selection and probabilistic classification methods and compare them to the methods presented in [7] on cfMeDIP-seq data across varying sequencing depths and cancer types.

### Description of materials

The cfMeDIP-seq data set used in this work was received from the authors of [7] by request. We used preprocessed read count data, where the sequencing read counts have been determined for genomic windows of length 300bp. The details of the data processing can be found from [7]. The discovery and validation cohorts consist of 189 and 199 samples respectively. The number of samples in each of the eight classes (corresponding to the healthy controls and 7 cancer types) is presented in Table 1. As in [7], the features for the classifiers are selected from a set of 505027 genomic windows, which consists of CpG rich regions, CpG islands, shores and shelves and FANTOM5 enhancers.

**Table 1:** Number of samples in each class in discovery and validation cohorts.

| Class | Discovery cohort | Validation cohort |
| --- | --- | --- |
| Healthy controls (Normal) | 24 | 62 |
| Renal cancer (RCC) | 20 | - |
| Pancreatic ductal adenocarcinoma (PDAC) | 24 | 47 |
| Colorectal cancer (CRC) | 23 | - |
| Bladder cancer (BLCA) | 20 | - |
| Breast cancer (BRCA) | 25 | - |
| Lung cancer (LUC) | 25 | 55 |
| Acute myeloid leukemia (AML) | 28 | 35 |
| Total | 189 | 199 |

### Workflow

We followed the workflow of the feature selection, model training and evaluation of the models presented in [7] but some modifications and additions were done. The workflow is illustrated in Fig. 1. First, we subsampled both discovery and validation cohort data sets and generated 100 data splits of the discovery cohort. In each data split, the discovery data was divided with 80%-20% ratio into balanced training and test sets using `caret` R package [23]. We utilized the scripts from [7] for generating the data splits. For each data split, we selected features with different feature selection methods using the corresponding training data set, with a data transformation applied when applicable.

**Figure 1:**
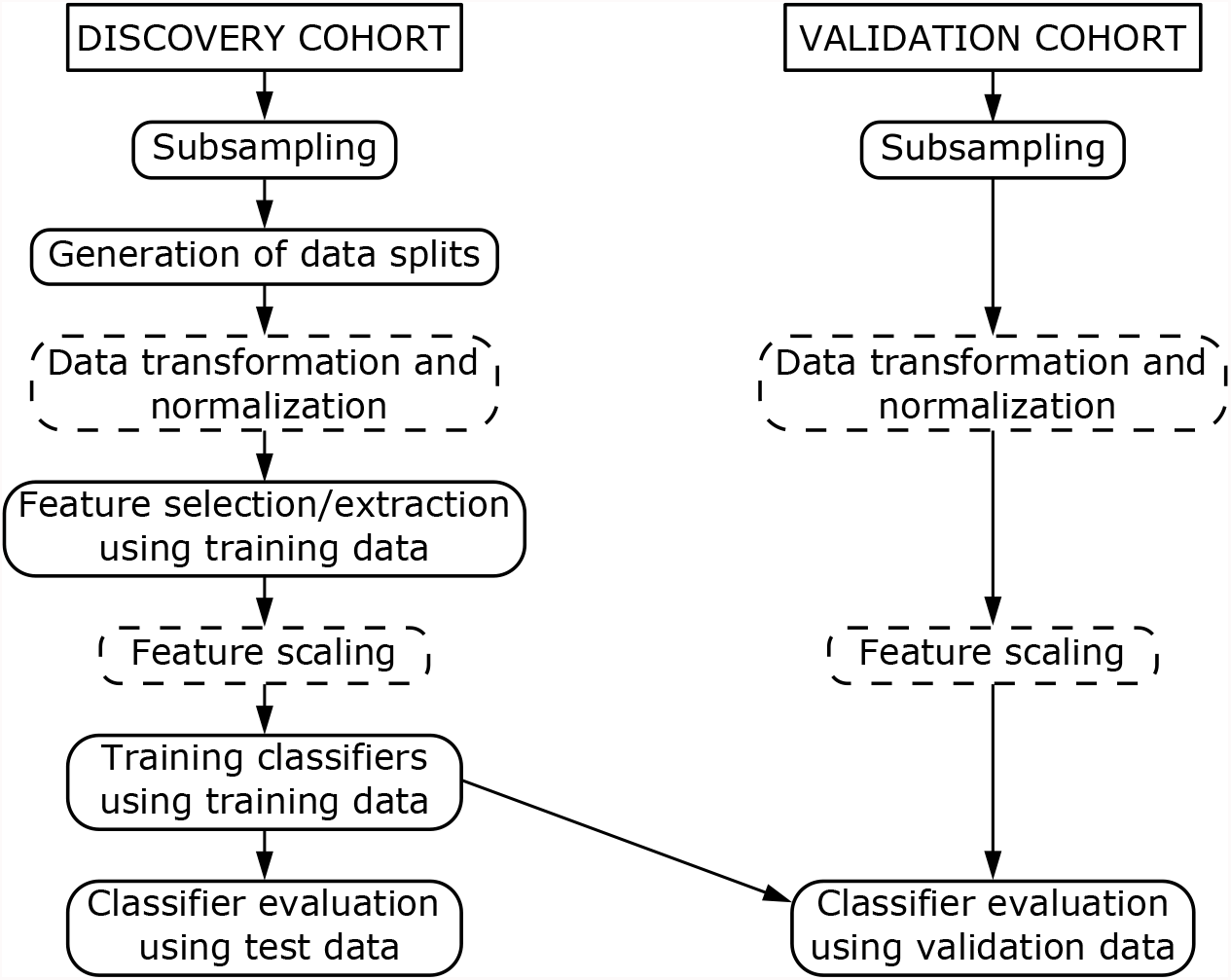
Workflow of the analysis. The boxes with dashed lines are steps that are performed when applicable. The validation data transformation, normalization and feature scaling are performed using scaling factors based on the training data. Each step after the generation of data splits is performed for each data split separately.

We trained the classifiers with the training data, using the found features. This resulted in 100 × 8 binary classifiers for each classification method. We made predictions for the corresponding test data sets to evaluate the classifier performances. Finally, we applied the trained models to the validation cohort and the classifier performances were again evaluated.

Our modifications and additions to the workflow of [7] include data sub-sampling, introduction of another version of the data transformation, addition of Fisher’s exact test for DMR finding and PCA and ISPCA methods for feature extraction, training the original classifier method with different types of features and addition of two different logistic regression classifiers with sparsity promoting priors.

### Data preprocessing

#### Data subsampling

The read count data was subsampled to simulate a lower sequencing depth than in the original data. The total read count in the discovery cohort data set, calculated from the preprocessed read count data, varies between 10659729 and 67228099 before extracting the 505027 genomic windows of interest. The thinning was carried out by randomly sampling the desired number of reads without replacement from all genomic windows. All reads had equal probability of being picked, and the probability of getting a read from a genomic window was proportional to the read count of the window in question. The subsampling was performed for all samples separately. This way we generated three subsampled versions of the data set with 10^4^, 10^5^ and 10^6^ reads per sample. The highest value, 10^6^, is already a magnitude lower than in the original, non-thinned data. After the thinning the 505027 genomic windows could be extracted.

#### Data transformation

Depending on the classification model, the count data can be used as it is or transformed. We used logarithmic transformations as proposed in [7]

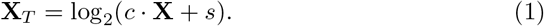

The transformation used in [7] is obtained with *c* = 0.3 and *s* = 10^−6^, but we also experimented with a modified version where *s* = 0.5. Transformation was applied to both discovery and validation cohorts when applicable. The difference of these transformations is best visible for the zero-count genomic windows. The original transformation maps the zero counts far away from the nonzero counts, while with the modified version the gap between the zero counts and nonzero counts is more moderate.

#### Scaling and normalization

Before fitting probabilistic classification methods, the count data was normalized based on the total read count in the 505027 genomic windows. This was done by dividing the read counts of these genomic windows with the sum of the read counts and multiplying with the mean of the read count sums over the discovery cohort. The read count normalization accounts for possible differences in sequencing depth between samples. In the case of the subsampled data, where the total read counts per sample were equal, this step should have affected only little. However, this is not the case with the non-thinned data where the total read counts varied greatly. Furthermore, the features were standardized to have zero mean and standard deviation of 1, so that the same prior mean and scale can be used for all features during the probabilistic classification step. The scaling factors were calculated based on the training data of each data split. Validation cohort was normalized and scaled accordingly, using normalizing and scaling factors of the training data.

### Methods for feature selection

#### Moderated t-statistic

To generate results with the same methods as in [7], we used the same DMR finding method. Moderated t-statistic implementation from the `limma` R-package [24] was applied to pick 150 hypo- and hypermethylated DMRs for each one-vs-one comparison, totaling in 7 × 300 = 2100 DMRs per class. The 2100 DMRs are necessarily not unique, so the number of unique DMRs can be lower. This was repeated for each class and each 100 data split. DMR sets were produced with both of the data transformations described above.

#### Fisher’s exact test

Fisher’s exact test was performed to the count data as an alternative method to the moderated t-statistic based DMR finding. For each genomic window, a contingency table with one-vs-one comparison setting was formed of the training data and p-value was calculated with `fisher.test` function from the R package `stats` [25]. Genomic windows with zero counts for all samples in the training data set were removed before conducting the tests. The DMR finding over classes and data splits was performed in similar manner as in the case of moderated t-test described above.

#### PCA

We utilized the 2100-dimensional^1^ set of DMRs as the input for principal component analysis (PCA). The resulting projections of the DMR vectors on the principal components are then given as features for the binary classifiers. PCA was conducted for each data split and each class separately, using `prcomp` function from R package `stats` [25]. The data was shifted to be zero-mean and scaled to have unit variance by using `center` and `scale` options of the `prcomp` function. The found components were standardized by dividing them with their standard deviations. Test and validation data was normalized and scaled with scaling factors calculated with the training data.

#### ISPCA

Iterative supervised principal component analysis (ISPCA) [20] is a method for finding features that are most relevant for predicting the target value. Following the notation of the description of the algorithm in [20], let us call the matrix of size *N*_samples_ ×*N*_features_ containing the original features as **X** and the target value vector of length *N*_samples_ as **y**. The process of finding supervised components iterates three steps. First, scores *S*(**x**_*j*_, **y**), *j* = 1, …, *N*_features_ which tell how relevant each feature **x**_*j*_ is for predicting target value **y** are calculated and the features that have a score higher than threshold *γ* are chosen. From these features, **X**_*γ*_, the first principal component **v**_*γ*_ is calculated. The threshold *γ* should be chosen so that the score *S*(**z**_*γ*_, **y**), where **z**_*γ*_ is the projection of **X**_*γ*_ onto the principal component, is maximized. Finally, the variation explained by the found feature is substracted from **X**, so that a modified feature matrix **X**′ is retrieved. **X**′ is then used as the starting point for the next iteration.

For our case with eight classes, both binary and multiclass approaches are possible. In the multiclass setting, features maximizing the score for one class versus other classes are searched for each class separately, and the feature with maximum score is picked. That is, the new component separates one of the classes from the others the best. With the multiclass approach ISPCA needs to be run only once per data split. For the binary approach, the multiclass labels have to be transformed into one-vs-rest binary labels before running ISPCA. ISPCA is then run for each class separately.

As finding too many supervised components might lead to overfitting, the ISPCA method includes a permutation test approach to calculate the p-value of there being relevant information left in **X**′. This test can be conducted after each iteration of finding supervised components and when the p-value exceeds a desired threshold, no more supervised components are searched. Non-supervised PCA can then be performed for **X**′ to retrieve up to *N*_samples_ − 1 components in total. We used the default threshold for the p-value, which is 0.1.

We used the ISPCA implemention from R package `dimreduce` [26], namely function `ispca`. We gave the read counts for all 505027 genomic windows as input to the analysis. Before running ISPCA, the data was normalized for total read counts and transformed with the new version of the data transformation. We used the `center` and `scale` options of the `ispca` function to standardize the input data and the option `normalize` to scale the returned components. Test and validation data were standardized and scaled accordingly.

### Classification methods

#### GLMnet

GLMnet classifiers were trained and evaluated as described in [7] by utilizing the provided R scripts. Briefly, the model training utilities from R package `glmnet` [9] were employed to learn a binomial one-vs-rest GLMnet classifier with the found DMRs as model features. The binomial GLMnet model corresponds to the logistic regression with elastic net regularization on the model coefficients [9]. The model is fitted by maximizing the penalized log-likelihood

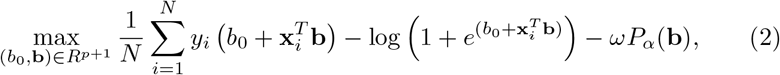

where *y*_*i*_ is the binary response variable, **x**_*i*_ is the corresponding feature vector, *N* is the number of observations, *b*_0_ and **b** are the intercept and coefficient parameters of the model and *P*_*α*_(**b**) is the penalty term multiplied with penalty parameter *ω*. Elastic net penalty is the sum of ridge-regression and Lasso penalties

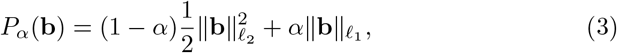

where mixing parameter *α* controls the proportions of the two penalty terms. If *α* = 0, elastic net simplifies into ridge regression and if *α* = 1 the penalty term becomes the same as in Lasso. As in [7], the parameters *ω* and *α* were optimized using three iterations of 10-fold cross validation for grid values *ω* = {0, 0.01, 0.02, 0.03, 0.04, 0.05} and *α* = {0, 0.2, 0.5, 0.8, 1}. Before training the binary classifiers, data transformation was applied. We also trained GLMNet classifiers with the DMRs found with Fisher’s exact test and with the moderated t-test using the modified version of the transformation.

#### Logistic regression with regularized horseshoe prior

In logistic regression model, each of the elements in the target vector **y** = (*y*_1_, …, *y*_*N*_) containing binary outcomes is assumed Bernoulli distributed with parameter *p*_*i*_, *i* = 1,.., *N*, where *N* is the number of samples. A linear model can then be connected to parameter *p*_*i*_ with inverse-logit function

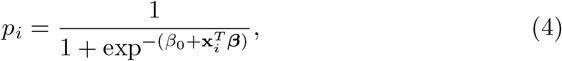

where *β*_0_ is an intercept term, ***β*** is a coefficient vector and **x**_*i*_ is a vector containing the values for the chosen features for sample *i*. After estimating the model coefficients, the class of a new sample can be predicted by calculating 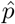 and using 0.5 (or some other value) as decision boundary. To classify the eight different classes, we trained logistic regression models for each class separately by first binarizing the class labels using one-vs-rest approach.

The regularized horseshoe prior [21] is a technique to achieve sparsity in a regression model when the number of features is large and only few of them are expected to be relevant, and thus should have nonzero regression coefficient. The regularized horseshoe prior enforces sparsity to the regression coefficients by defining the scale of the coefficients to be a product of local and global terms, where the global term pulls all coefficients towards zero, while the local term allows the relevant features to have nonzero coefficients. The prior for the regression coefficients *β*_*j*_ can be expressed more formally as

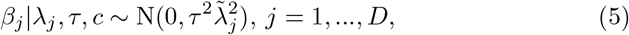

where *D* is the number of features and the modified local scale parameter 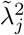 is defined as

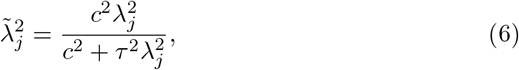

where the local scale parameter *λ*_*j*_ is given a half-Cauchy prior

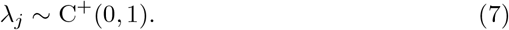

The modified local scale parameter makes sure that all coefficients are shrunken at least a little, including the relevant coefficients as well. The parameter *c* controls the magnitude of the largest coefficients and is given an inverse-Gamma prior

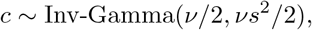

where *ν* and *s* were set to 4 and 2 respectively, following the default values of the regularized horseshoe prior implementation in R package `brms` [27]. The global scale parameter *τ* has also a half-Cauchy prior

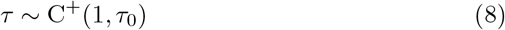

with scale *τ*_0_ defined as

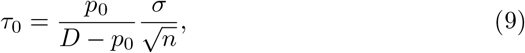

where *D* is the number of features, *n* is the number of training samples, *p*_0_ is the expected number of nonzero coefficients and *σ* is a pseudo standard deviation. The RHS prior enables using the knowledge on the expected number of nonzero coefficients to define the global scale parameter hyperprior. We set *p*_0_ = 300 after experimenting with a few different options and comparing the resulting coefficient posteriors. The pseudo variance for a model with binomial data and logit link function proposed in [21] is

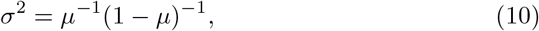

where *μ* is replaced with sample mean 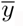. The intercept term *β*_0_ is handled separately and is given a Gaussian prior with mean 0 and standard deviation of 10.

The logistic regression model with horseshoe prior was fitted with different kinds of features: moderated t-statistic DMRs with the original data transformation, the moderated t-statistic DMRs with the modified data transformation, Fisher’s exact test DMRs, PCA coordinates and ISPCA coordinates. The original data transformation was applied before model fitting on the corresponding moderated t-statistic DMRs and Fisher’s exact test DMRs, while the modified data transformation was applied on the corresponding moderated statistic DMRs. Before logistic regression we normalized the features for total read counts and standardized each feature to have zero mean and variance of 1.

The model was implemented with the R version of the probabilistic programming language Stan [28]. For the logistic regression with horseshoe prior, we adopted the example code presented in [21], where model parametrization proposed in [29] was used.

The predictions for test and validation data sets were made by calculating the posterior predictive probabilities. For the *i*^th^ test/validation sample 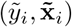 we compute an approximation of the probability of it belonging to the class of interest 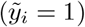 using posterior samples of *β*_0_ and ***β***

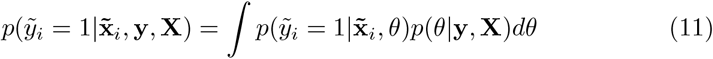

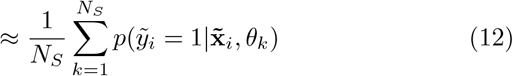

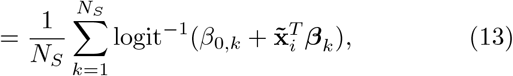

where *N*_*S*_ is the number of MCMC samples retrieved from the posterior distribution and *θ*_*k*_ = (*β*_0,*k*_, ***β***_*k*_) denotes the *k*^th^ parameter sample. See description of the MCMC sampling method below.

#### Logistic regression with DMR count variables

We also formulated a simpler logistic regression model for binary classification, where the model includes only an intercept term and two features. These two features are the numbers of hypermethylated and hypomethylated DMRs with nonzero read counts. The DMRs can be found either with the moderated t-statistic or Fisher’s exact test. The two features were normalized based on total read counts. and scaled to have zero mean and standard deviation of 0.5. The model intercept and coefficients were given Cauchy priors with scale parameters 10 and 2.5, respectively, as recommended in [30]. The model was implemented with Stan. The classification of the test and validation sets are done in the same way as for the logistic regression model with RHS prior.

### Sampling from posterior distributions with Stan

Stan uses MCMC sampling to retrieve samples from the posterior distribution, specifically the no-U-turn sampler (NUTS) algorithm, which is a variant of the Hamiltonian Monte Carlo algorithm [22]. The parameters for sampling the user can define the number of MCMC chains, number of samples per chain, maximum tree depth and target acceptance rate. The sampling parameters for our models are presented in Table 2.

**Table 2:** Sampling parameters for the models implemented with Stan. Values marked with * are the default values in Stan.

| Sampling parameter | LR RHS | LR RHS<br>ISPCA/PCA | LR DMR<br>counts |
| --- | --- | --- | --- |
| Number of chains | 4* | 4* | 4* |
| Number of samples per chain | 2000* | 4000 | 3500 |
| Max. tree depth | 10* | 10* | 10* |
| Target acceptance rate | 0.8* | 0.99 | 0.9 |

### Evaluation of the classification methods

The evaluation of the classification methods was performed in similar manner as in [7] and the distributed code for the publication was utilized when implementing the methods. For each method, each of the eight binary classifiers for each data split were used to classify the corresponding test data set and the validation data. The performance of the classifiers was evaluated by calculating class-specific area under receiver-operating characteristics curve (AUROC) and area under precision-recall curve (AUPRC) statistics. The distribution of the statistics for each class over data splits can then be described by calculating median and quantile statistics or by plotting boxplots. The AUROC statistics are presented as barplots and scatterplots. In addition, median AUROC and AUPRC values are presented as tables. For the validation cohort, we also calculated a mean of the class predictions over the data splits and plotted a receiver-operating characteristics curve (ROC) and calculated corresponding AUROC and AUPRC values for each of the four classes in the cohort.

### Evaluation of the methods with an independent data set

To see whether the results produced with the data set from [7] generalize to other data sets, we ran the same analysis (excluding data subsampling) for an independent intracranial tumors data set by [11]. The data was downloaded as reads per kilobase per million mapped reads (RPKM) values for each genomic window from [31]. The data set consisted of 161 samples of 6 different tumor types. Table 3 shows the number of samples in each class. As the data was given as RPKM values, there was no need to perform normalization with respect to total read counts.

**Table 3:**
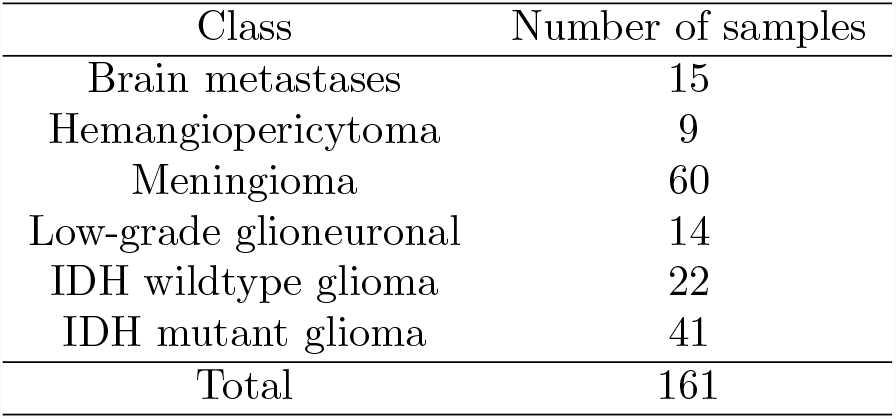
Number of samples in each class in the intracranial tumors data set.

## Results

### Feature selection

#### Comparison of the DMR finding methods

We tested three different methods to find differentially methylated regions that can be utilized as features in the classification: the moderated t-test method used in [7], moderated t-test with new data transformation and Fisher’s exact test. To compare the DMRs found with the different methods, we plotted Venn diagrams to find their overlaps. In Fig. 2, for each method, all of the DMRs for the 100 data splits were first combined and duplicates were removed to keep each DMR in the set only once. This was done for each class separately. In Supplementary Fig. S1, the Venn diagrams were generated from DMRs that were found in 50 or more data splits of the total 100.

**Figure 2:**
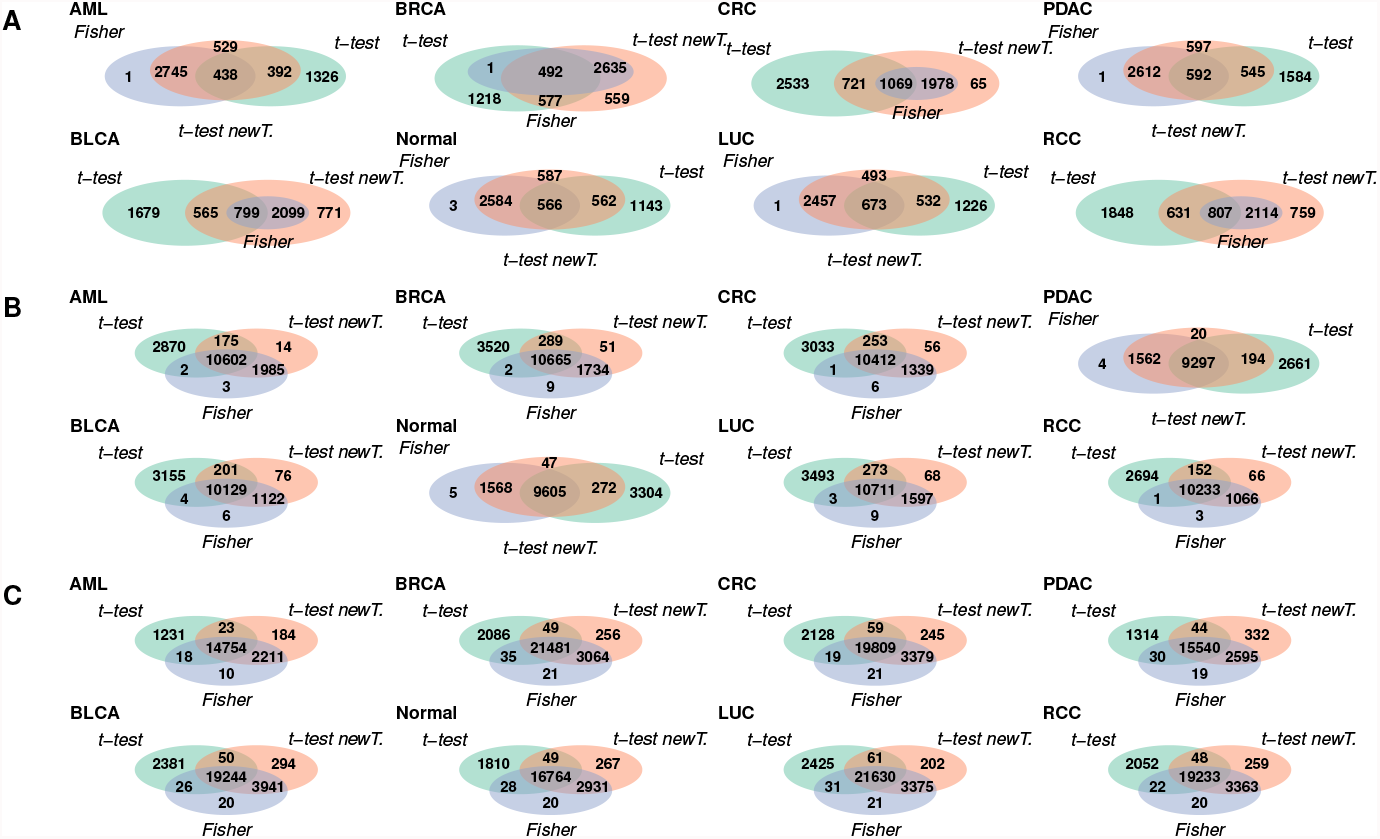
Number of overlapping DMRs between the DMRs found with Fisher’s exact test, moderated t-test with the original data transformation and moderated t-test with new data transformation. DMRs from the 100 data splits were combined before the comparison. The results are presented for thinning with total read counts 10^4^, 10^5^ and 10^6^ in figures **A, B** and **C**, respectively.

Comparing Fig. 2 and Supplementary Fig. S1, we can see that the number of DMRs is overall higher in Fig. 2, where the DMRs were not filtered. This suggests, that many of the DMRs are only found in less than half of the data splits. In Fig. 2 we can also notice that overall the number of DMRs is smaller when the total read count is smaller, i.e. data has been thinned more. This might mean, that when the total read count is smaller, the found DMRs are more consistent between different data splits, resulting in a smaller number of DMRs. Supplementary Fig. S1 supports this, as the number of filtered DMRs is overall higher in Supplementary Fig. S1**A** than in Supplementary Fig. S1**B** and Fig. S1**C**. However, the overlap between all three methods is low in Supplementary Fig. S1**A**, indicating the different DMR finding methods worked rather inconsistently in the case where the total read count was low.

In most cases a large fraction of the DMRs was shared with all of the three methods. The overlap between the Fisher’s exact test and t-test with the new data transformation was often quite high, as was the number of DMRs unique to the original t-test method. The numbers of DMRs unique to either Fisher’s exact test or the t-test with the new data transformation were often low in comparison. The overlaps between the original t-test method and the two other methods separately were quite modest compared to the overlap between Fisher’s exact test and t-test with new transformation. Altogether, it seems that a large part of the DMRs was shared between all of the methods, which suggests that DMRs can indeed be identified reliably from cfMeDIP-seq data. On the other hand, there were also DMRs not shared by all three methods. These DMRs may partly explain the differences in the performances of the classifiers utilizing these DMR sets.

Lists of DMRs for each of the three DMR finding methods are available in Supplementary files 2-4. See Section Availability of data and materials for details.

#### Comparison to the RRBS-seq based DMCs

In [7], DMRs identified from cfMeDIP-seq data were compared to differentially methylated cytosines (DMCs) identified from reduced representation bisulfite sequencing (RRBS-seq) data that was obtained from solid samples. Shen et al. (2018) presented two sets of RRBS-seq DMCs: from comparisons between normal and tumor tissues as well as between tumor tissue and normal peripheral blood mononuclear cells (PBMCs). The comparison of cfMeDIP-seq DMRs and RRBS-seq DMCs presented in [7] showed that there was significant enrichment in concordantly hypermethylated and hypomethylated cfMeDIP-seq DMR and RRBS-seq DMC pairs. To see if there was overlap still after data subsampling, we made a simple comparison between the DMRs found from the thinned cfMeDIP-seq data and the two types of DMCs provided in [7]. The comparison was carried out by finding the cfMeDIP-seq DMRs from PDAC class vs. normal class comparison which had one or more overlapping DMCs. The direction of differential methylation was required to be the same in the RRBS-seq DMCs and cfMeDIP-seq DMRs when finding the overlaps. The number of such cfMeDIP-seq DMRs was calculated for each data split for each of the three subsampling versions. This was repeated for the three DMR finding methods.

The results of the comparison in Fig. 3 show that the number of cfMeDIP-seq DMRs with overlapping DMCs in a data split was overall quite low, ranging from 0 to 14. The overlap with the DMCs from the comparison of tumor tissue and normal tissue was generally lower than with the DMCs from the comparison of tumor tissue and PBMC. The severity of the thinning does not seem to affect the number of cfMeDIP-seq DMRs with RRBS DMC overlap, suggesting that the most significant DMRs can still be identified from the subsampled data as well. Overall the level of overlaps with the RRBS-seq DMCs seemed to be the same for all three DMR finding methods, with the exception of the thinning with total read count 10^4^, where the moderated t-test with new data transformation had considerably more DMRs with DMC overlap than the two other methods.

**Figure 3:**
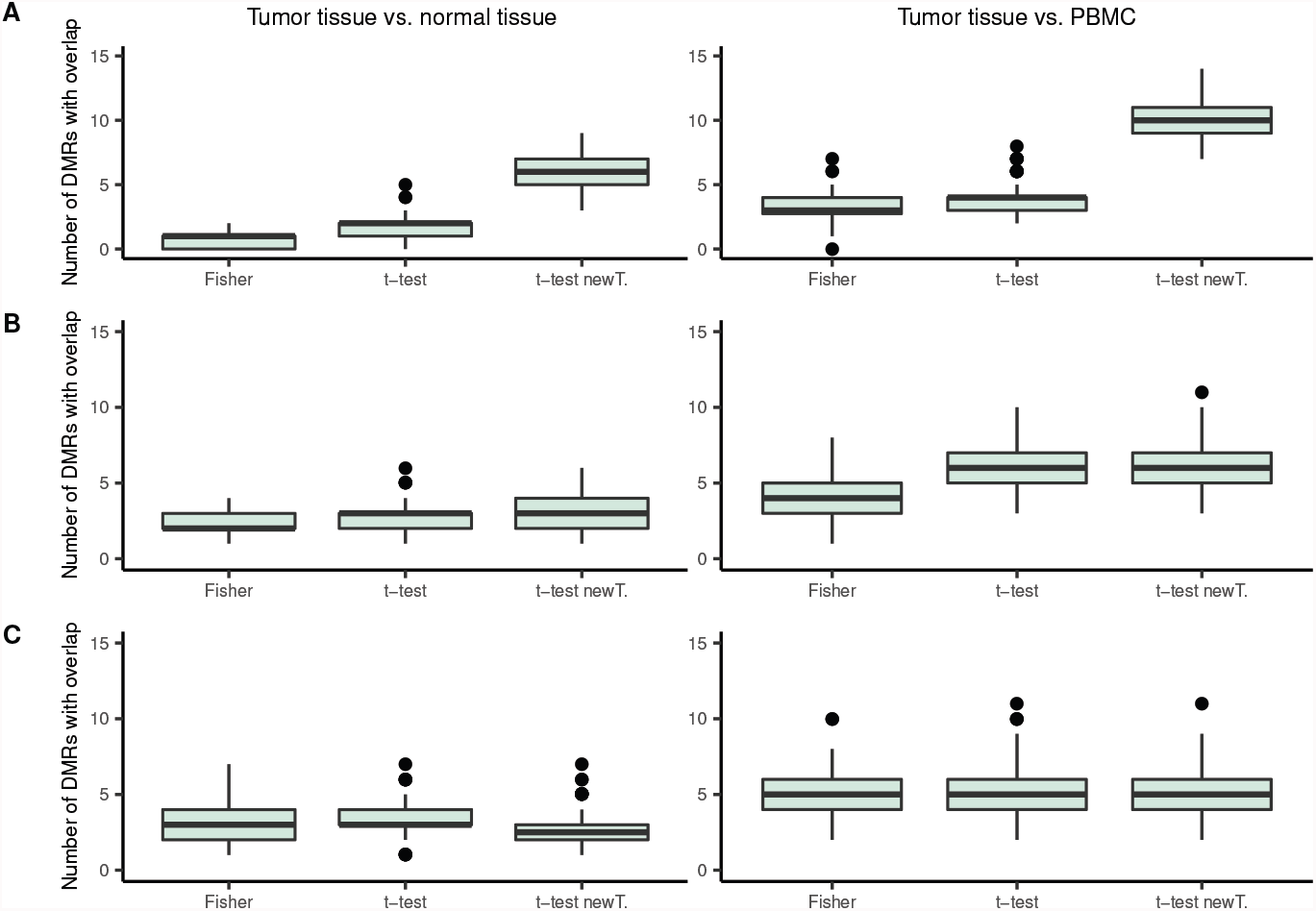
Boxplots of the number of cfMeDIP-seq DMRs having overlap with RRBS-seq DMCs in one data split, for each of the DMR finding approaches. The left side column shows the results for the RRBS-seq DMCs from comparing PDAC tumor tissue to normal tissue, while the right side column shows the results for RRBS-seq DMCs from comparing PDAC tumor tissue to PBMC. The results are presented for thinning with total read counts 10^4^, 10^5^ and 10^6^ in figures **A, B** and **C**, respectively.

#### PCA and ISPCA

Fig. 4 demonstrates the retrieved features from the three different dimension reduction approaches, PCA, binary ISPCA and multiclass ISPCA, for one of the data splits in the case of the subsampled data with total read count 10^6^.

**Figure 4:**
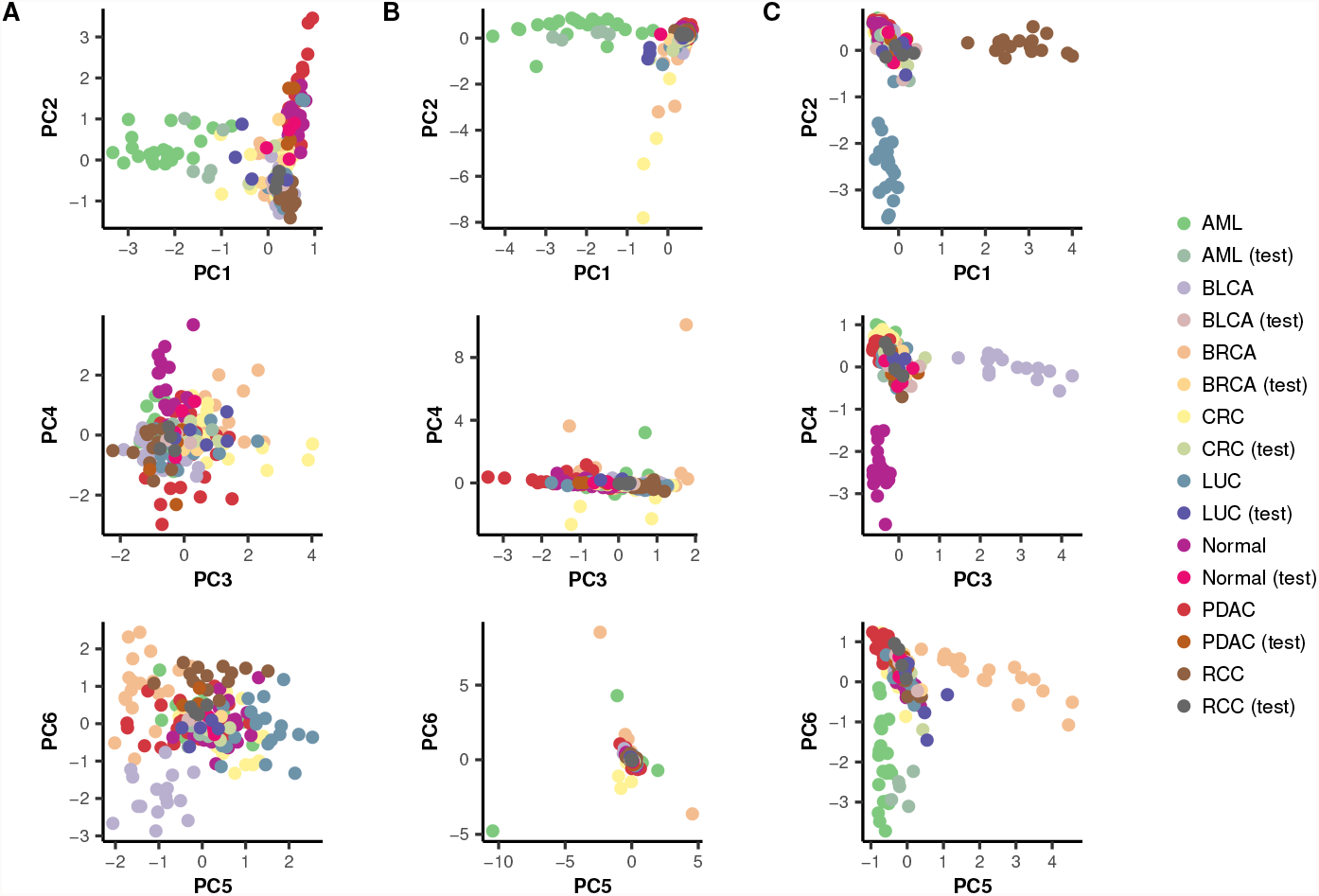
An example of the six first principal components from PCA and two ISPCA approaches for one of the data splits. The training and test samples from the discovery cohort were plotted with different colors to demonstrate how well the test samples will blend in with the training samples. **A**: Components from PCA, using AML class DMRs as input. **B**: Components from binary ISPCA, where class labels were set to 1 for the samples in AML class and to 0 for the samples from other classes. **C**: Components from multiclass ISPCA.

Fig. 4**A** shows the first six principal components from the standard PCA, two at a time. The input data to the analysis was the DMR set for the AML class, and based on the plot, the AML class could be separated well from the other classes using PC1. The training and test set samples from AML class seemed to mix to some extend, which indicates that the principal component generalized well to the test samples. The other plotted components seemed not to separate AML class from the others.

We generated similar plots for the binary and multiclass ISPCA approaches, presented in Fig. 4**B** and Fig. 4**C**, respectively. For the binary ISPCA we set the samples from the AML class to have label 1 and for the other classes the label was set to 0. We gave data from all 505027 genomic windows as input to ISPCA. This resulted again in the first component separating the AML class from the other classes quite well, while the test samples blended in with the training samples. The rest of the plotted principal components did not separate the AML class from the other classes. The first six components from multiclass ISPCA seemed to each separate one class from the others. The only class for which the test samples blended in with the plotted training samples was AML, which could indicate that making good predictions with a model that uses these components as features could be hard.

With the two other subsampled data versions, with total read counts 10^4^ and 10^5^, the ISPCA approaches occasionally ended up in a situation where no supervised components were found. In the case of more heavily thinned data, there seemed sometimes not to be enough information to find any supervised components, and the method returned standard PCA components only. The ISPCA methods could not always separate the classes well, which might be because we gave all 505027 genomic windows as input to them. On the other hand, the standard PCA approach which utilized the class-specific DMRs only did not thrive in such a situation either.

#### Experiments with different DMR numbers

To justify the choice of picking 300 DMRs per one-vs-one class comparison as features for the classifiers, we performed a comparison of GLMnet classifiers with different DMR numbers. The tested DMR numbers were 100, 300 and 400. The AUROC and AUPRC values for each class are presented in Supplementary Fig. S6-9. The classifiers were evaluated on both discovery and validation cohorts. The figures show that the classifiers performed approximately equally well with all DMR numbers. The classifiers with DMR number 100 have a slightly weaker performance, but there are no big differences between 300 and 400. The differences in AUPRC values are greater than in AUROC values. We compared DMRs from both Fisher’s exact test and moderated t-test (with new data transformation), and they both gave similar results. The results are as expected, as choosing the DMRs to be used as features works as a preselection, and the regularization of the GLMnet model finally picks the most important features. Based on these results, 100 DMRs might be too few. The benefit of increasing the number over 300 seems to be rather small, so we decided to proceed with 300 DMRs as proposed in [7].

#### Discovery cohort

After finding the model features, the different classification models were fitted using the training data sets of the discovery cohort. The data was partitioned to training and test data sets 100 times, and each of the trained models was evaluated with corresponding test set. This resulted in 100 AUROC values per each of the eight classes, from which we can calculate median AUROC values which are presented in Supplementary Fig. S2. Based on this figure, the overall trend was that the lower the total read count is after the data subsampling, the lower the median AUROC values were. This is expected, as the more the data is thinned the less there is information for us to use for the classification task. When the total read count was 10^4^, the median AUROC levels were very similar for all methods and for all classes. When the total read count was higher, some classes such as AML and PDAC began to stand out with higher median AUROC values, while some classes such as BRCA, CRC and LUC had lower median AUROC values even with higher total read counts.

To better compare each of the methods to the original approach, we computed the differences between median AUROCs (Fig. 5). The differences were calculated for each class separately, but a mean over classes is also presented for overall performance comparison purposes. Looking at the means of the AUROC median differences over classes, all GLMnet-based approaches seemed to perform overall equally well. However, there was some variance in the class-specific median AUROC differences for the GLMnet with Fisher’s exact test DMRs as features when the subsampled data had total read count 10^4^ or 10^5^. Similarly, the two logistic regression models with RHS prior using the original moderated t-test DMRs or the moderated t-test with new data transformation DMRs seemed to work equally as well as the original GLMnet method in all of the three thinning versions. The LR model with RHS prior and Fisher’s exact test DMRs, on the other hand, had a positive mean AUROC difference value when the sequencing depth was low. The logistic regression models with DMR count variables seemed to perform overall as well as the original GLMnet method or slightly worse.

**Figure 5:**
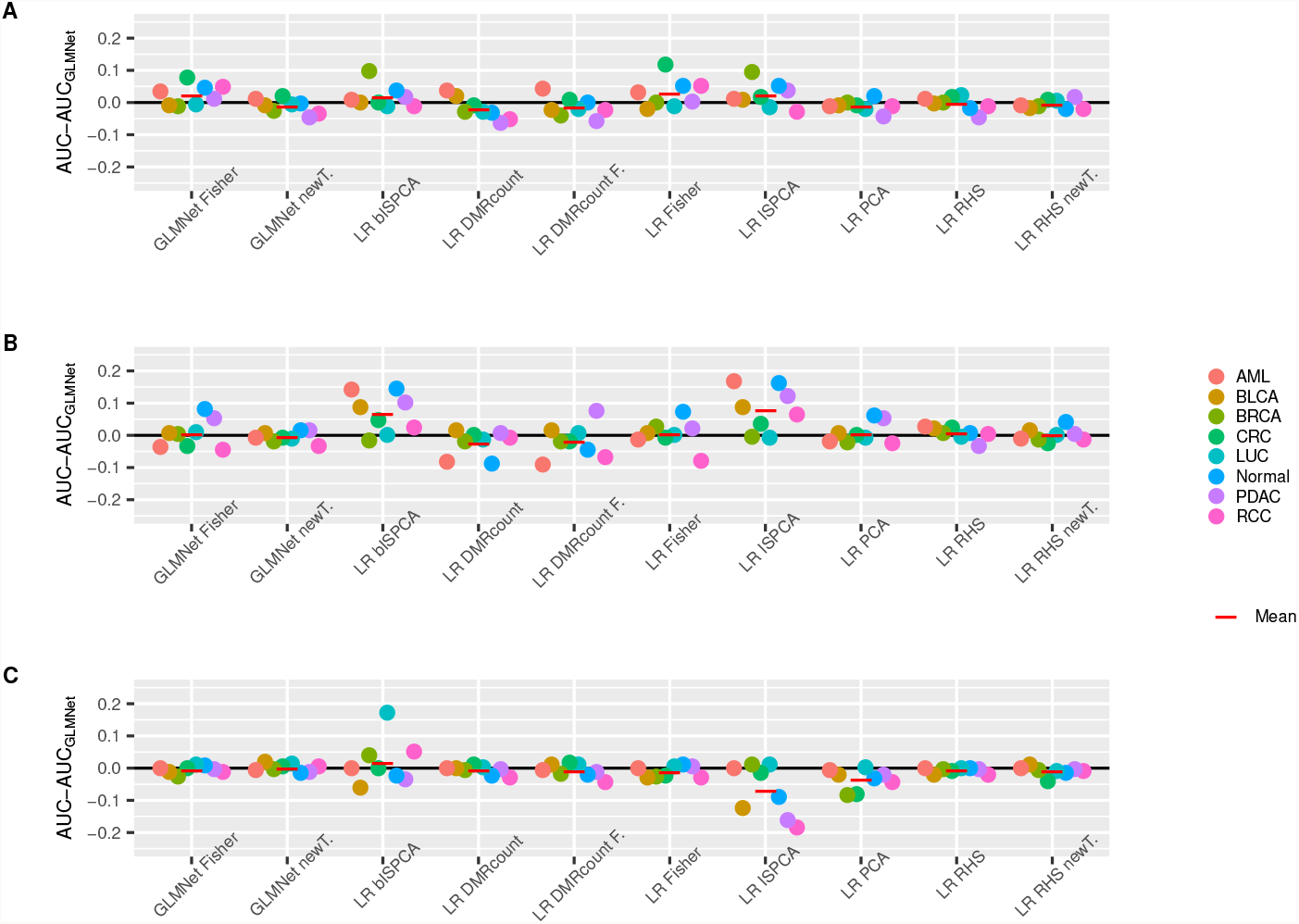
The differences between the test set AUROC medians over the 100 data splits for the original GLMnet method and other approaches. The AUROC was calculated for the test data sets in the discovery cohort. The results are presented for each class separately. Positive values indicate, that the compared (new) method had higher AUROC median than the original method and negative values indicate that the original method had higher AUROC median. The results are presented for thinning with total read counts 10^4^, 10^5^ and 10^6^ in figures **A, B** and **C**, respectively. The red lines indicate the means over the eight classes.

Continuing the comparisons with the LR models that use dimension reduction features, the logistic regression model with RHS prior that used PCA components as features worked approximately equally well as the original GLMnet method for all subsampled data versions. But perhaps the most promising methods in this comparison were the logistic regression models with RHS priors that used ISPCA components as features. Both the models using multiclass ISPCA and binary ISPCA components got higher median AUROC values than the original GLMnet method for most of the classes in Fig. 5 **A** and **B**. Surprisingly, when the total read count after thinning was set to 10^6^ (Fig. 5 **C**), the model using multiclass ISPCA components performed overall considerably worse than the original GLMnet method. However, the mean median AUROC differences over all classes was still slightly on the positive side for the model using binary ISPCA components.

Additionally, median AUROC and AUPRC values over the data splits are presented in Supplementary Tables. S1-S3. The ranking of the methods was approximately the same based on either AUROC or AUPRC values. However, there was more variation in the AUPRC values. Overall, the AUPRC values were much lower than AUROC values as expected, but the values tended to increase as the simulated sequencing depth increased in the same manner as AUROC values did.

#### Validation cohort

The models trained with the discovery cohort training sets were used to predict the class of samples in validation data set. The validation data was also thinned in the same manner as the discovery cohort data. The predictions were made with the models for each of the 100 data splits, and AUROC values were calculated for each set of predictions.

The AUROC medians over the 100 values were calculated and they can be found from Supplementary Fig. S3. As for the discovery cohort in Supplementary Fig. S2, the median AUROC values seemed to be overall higher when the total read count is higher. When compared to the discovery cohort results, there seemed to be more variance in the performance of different methods for the validation data.

We also calculated the median AUROC differences between the original GLMnet method and the new methods (Fig. 6). The logistic regression models with RHS prior that used the components from multiclass or binary ISPCA did well in the discovery cohort performance comparison. Conversely, these two models did not perform very well with the validation data and at best performed equally well as the original GLMnet method. Logistic regression with RHS prior using PCA components seemed to do slightly better than the ISPCA approaches, as the mean of median AUROC differences reached positive value for the case with total read count 10^5^. The overall performance of the logistic regression models with DMR count variables was usually as good or slightly weaker than for the original approach. However, for the subsampled data with total read count 10^4^, the approach using Fisher’s exact test DMRs had a considerably positive mean difference value. The GLMnet methods with the alternative DMR choices both performed approximately equally well as the original GLMnet method with all of the three subsampling versions.

**Figure 6:**
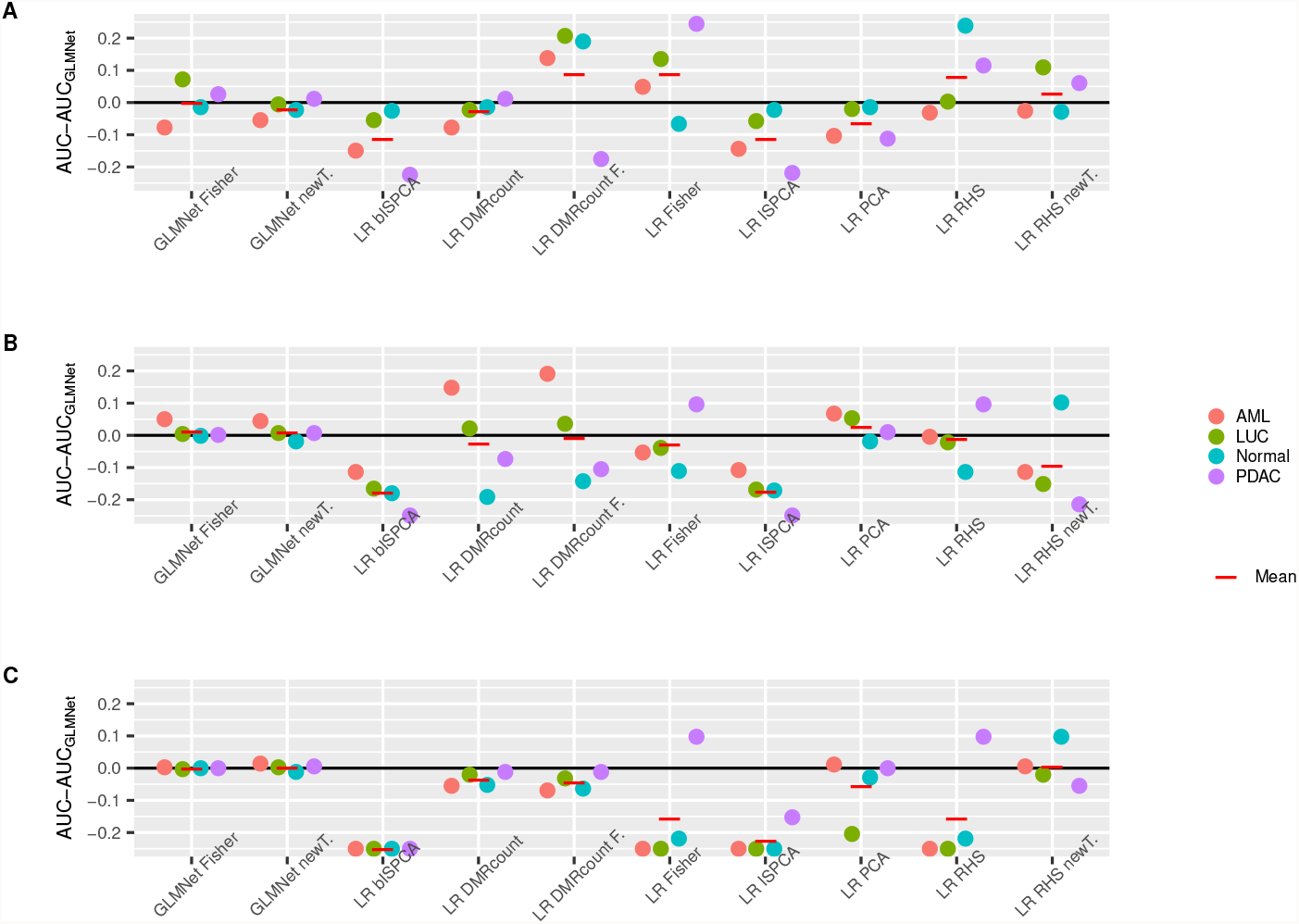
The differences between the validation cohort AUROC medians over the 100 data splits for the original GLMnet method and other approaches. The results are presented for each class separately. Positive values indicate, that the compared method had higher AUROC median than the original method and negative values indicate that the original method had lower AUROC median. The results are presented for thinning with total read counts 10^4^, 10^5^ and 10^6^ in figures **A, B** and **C**, respectively. The negative values were truncated to − 0.25. The red lines indicate the means over the four classes.

All in all, compared to the discovery cohort results, there seemed to be a lot more variation in the AUROC differences for the validation data set. When the total read count was 10^6^, i.e. closest to the original read counts, none of the methods had positive mean difference over the classes. But when the thinning became more severe, there were some cases where other methods outperformed the original method.

As in [7], we also calculated the mean of the validation set predictions over the 100 data splits and produced ROC curves for each class. In Fig. 7, we can see a general trend of the area under the curves getting higher the higher the total read count is. The ROC curves confirm the findings we did based on the median AUROC differences. When the total read count was 10^6^, several methods performed approximately equally well as the original GLMnet method, but rarely outperformed it. When the total read count was 10^5^ or 10^4^, some methods had their ROC curves surmounting the original method’s curve for some of the classes.

**Figure 7:**
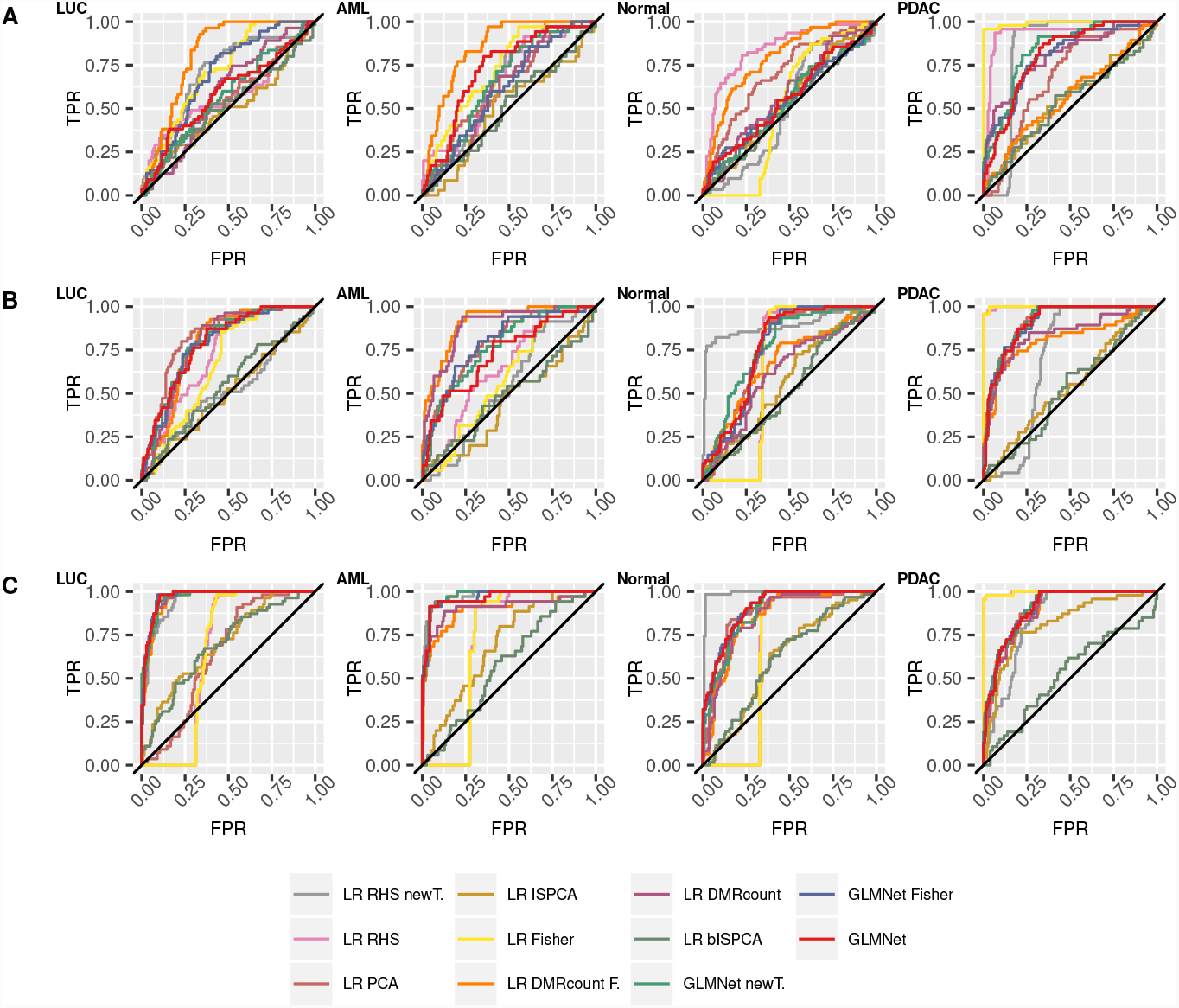
Validation cohort ROCs calculated with prediction means over 100 data splits and corresponding fitted models. The results are presented for each class separately. The results are presented for thinning with total read counts 10^4^, 10^5^ and 10^6^ in figures **A, B** and **C**, respectively.

Again, the corresponding median AUROC and AUPRC values are presented in Supplementary Tables. S4-S6. The conclusions about the AUPRC values are the same as for the discovery cohort.

#### Non-thinned data

Until now, the results we have presented have been for the thinned data set. To assess how well the classifiers presented in this work perform with the original non-thinned data set, we ran the feature selection methods and classifier training to the data without thinning it first. We calculated the median AUROC values, the median AUROC differences between the original GLMnet method and the other classifiers for both the discovery and validation cohorts and produced ROC curves of the mean predictions for the validation cohort. The results are presented in Supplementary Figs. S4 and S5.

The median AUROC values over the 100 data splits are presented in Supplementary Fig. S4 for both discovery and validation cohorts. We can see that overall the AUROC medians were higher for the more deeply sequenced, non-subsampled data, both for the discovery and validation cohorts, than for the thinned data sets. The differences between the methods were moderate for the discovery cohort, but for the validation cohort there were greater differences. There were also differences between the classes, indicating that some classes were easier to distinguish from the other classes with the used methods. For example, considering the discovery cohort, all methods reached median AUROC values close to 1 for the AML class, while for LUC class the values were at best around 0.875.

In Supplementary Fig. S5**A** we present the median AUROC value differences between the original GLMnet method and the other methods for the discovery cohort. For most of the methods the mean differences were close to 0, meaning that overall there was no difference in the median AUROC values for the original GLMnet method and other approaches. However, the performance of logistic regression model with RHS prior and multiclass ISPCA components and logistic regression models with DMR count features, the mean difference values were negative.

Supplementary Fig. S5**B** presents the median AUROC differences between the original method and the other classifiers for the validation data set. Based on this comparison, the overall performance of the other classifiers was again in most cases equally good or even considerably weaker than for the original GLMnet method. There were few exceptions in the case of the PDAC class, for which the logistic regression models with RHS prior using either of the two moderated t-test or the Fisher’s exact test DMRs had all approximately 0.1 higher median AUROC value than the original method. Also, the modified GLMnet approaches both had approximately equally good results as the original method.

From the validation cohort ROC curves in Supplementary Fig. S5**C** we can see, that the original GLMnet method performed similarly as in [7], with almost equally good performance for LUC, AML and Normal classes and slightly weaker performance for PDAC class. The corresponding AUROC values in [7] were 0.971, 0.980, 0.969 and 0.918 for classes LUC, AML, Normal and PDAC respectively. The ROC curves for the other two GLMnet approaches behaved quite similarly. For the rest of the methods, the ROC curves support the findings based on the median AUROC differences in Supplementary Fig. S5**B**.

The conclusions described above are supported with median AUROC and AUPRC values for discovery cohort in Supplementary Table S7 and for validation cohort Supplementary Table S8. However, in the case of the discovery cohort, there were some classes (BRCA, CRC, LUC) where the methods with highest AUROC and AUPRC values were not the same. The method with best AUROC did still have one of the top AUPRC values, so the ranking of the methods was similar with both statistics.

#### Results for independent data set

We repeated the analysis with our workflow (excluding data subsampling and total read count normalization) using the intracranial tumors data set from [11], to see whether the results generalize to other data sets. AUROC barplots for the data set can be found Supplementary Fig. S10, while the AUROC and AUPRC values are presented in Supplementary Table S9.

Our results are in line with the classifier performances reported in [11]. The AUROC values presented in [11] were 0.89, 0.95, 0.93, 0.82 and 0.71 for meningioma, hemangiopericytoma, low-grade glioneuronal, IDH mutant glioma and IDH wildtype glioma classes, respectively (AUROC was not reported for brain metastases).

Based on the AUROC barplots in Supplementary Figure. S10, GLMnet and logistic regression with modified t-test (with either version of the data transformation) and logistic regression with PCA components as features seemed to perform the best overall. The logistic regression models with DMR count features had considerably weaker performance. Classifiers with Fisher’s exact test DMR features also had lower AUROC values when compared to the classifiers using modified t-test DMRs. The AUPRC values in Supplementary Tables. S7 support these findings. The AUROC-based ranking of the methods with the non-thinned cfMeDIP-seq data from [7], shown in Supplementary Fig. S4, is similar. However, the differences between the methods seemed to be bigger with the intracranial tumors data set than with the data set from [7].

## Discussion

There seems to be a lot of variation in how well the compared methods performed with different classes, thinning versions and also between discovery and validation cohorts. For example, the LR model with RHS prior using binary and multiclass ISPCA features seemed to perform better than the original method when looking at the discovery cohort results for the lowest sequencing depth. But looking at the validation cohort performance, these two approaches had considerably weaker median AUROC values than the original GLMnet method. One possible cause to the differences between the three subsampled data versions is that the subsampling is done by taking random subsets of the original data, and that could cause differences even if the probabilities of obtaining reads from each of the genomic window were the same for all three thinning versions. The differences between the discovery and validation cohort data sets are visible with the non-thinned data set too, and this indicates that there are perhaps differences between the two cohorts, making it harder for the classifiers to perform well. However, the GLMnet-based methods seemed to all cope quite well with the validation cohort, especially if the data is non-thinned or the data has not been thinned very much. It could also be, that not all DMR finding methods and classifiers are suited to all classes. It could be considered, that different classifiers would be used with different classes to obtain optimal performance.

The overlaps between the DMRs found with the three different approaches showed that there are indeed differences between the methods, even if many of the DMRs were shared between all methods. We noticed, that the Fisher’s exact test and the moderated t-test with new data transformation often shared DMRs which were not found by the original moderated t-test method, while the moderated t-test found DMRs that were not found by other methods. The numbers of unique DMRs to Fisher’s exact test or the moderated t-test with the new data transformation were often low in comparison. The DMRs that were not found by all methods could be a source of differences in the classification results.

The dimension reduction techniques, PCA and two versions of ISPCA, showed varying performance. While logistic regression models using ISPCA components did not perform well with the validation data cohort, their AUROC values were promising for the discovery cohorts when total read count was 10^4^ and 10^5^. We also tested using DMRs as input data for multiclass and binary ISPCA. The classification results with LR model and RHS prior using the resulting principal components as features showed stable performance, but these approaches did not outperform the original GLMnet method neither in the discovery or validation cohorts.

Both logistic regression with RHS prior and GLMnet methods implement the logistic regression model and enable sparsity of the feature coefficients. The difference between these two models is the prior enabling sparsity and how the model is fitted. In our approach we use the regularized horseshoe prior and GLMnet utilizes elastic net regularization. On the model fitting side, we utilized probabilistic programming language Stan to obtain posterior samples of the model, while GLMnet model is fitted with cyclical coordinate descent approach [9]. The GLMnet model fitting is very efficient, but while sampling with Stan requires more computational resources, we obtain samples that inform us of the whole posterior distribution. If we look at the performance of the LR RHS method with the moderated t-test DMRs as features, we notice that with the thinned data sets the performance is approximately equally good as for the original GLMnet method. The same applies to the LR RHS method with moderated t-test with new data transformation. For the validation cohort, there are both classes for which the performance is either considerably better or considerably weaker. Based on this comparison, our implementation of the logistic regression method with enabled model sparsity seems to have potential to give better results than GLMnet method, but it is perhaps not as stable as GLMnet. Combining promising feature selection methods such as ISPCA with the LR RHS model could further enhance the classification.

Based on the promising results in TCR analysis application [32], we expected the simple logistic regression model with two DMR count variables to perform well with the lowest sequencing coverage due to model robustness. The results with the thinned validation cohort with total read count 10^4^ somewhat support this hypothesis, but the discovery cohort results were not as impressive.

The results for the independent intracranial tumors data set seemed to support our findings with the non-thinned data set, indicating that these results could generalize to other data sets. However, the differences between methods were greater with the intracranial tumors data set than with the one from [7].

## Conclusions

We performed a method comparison to investigate if we the classification accuracy of the classifier based on cfMeDIP-seq data could be improved in a case where the sequencing depth of a cfMeDIP-seq experiments is low. To simulate lower sequencing depths, we thinned a cfMeDIP-seq data set by randomly sub-sampling the reads of each of the samples. We obtained three data sets with total read counts of 10^4^, 10^5^ and 10^6^. Then we tested three different DMR finding methods and three different dimension reduction methods to find the features to be used in the classification. After finding the features, three different types of classifiers were trained, using the found features and performance was evaluated for discovery and validation cohorts. These steps were also performed for the non-thinned data set and an independent intracranial tumors data set.

Based on the comparisons between the performances of the classification methods, there seemed to be no one method that would consistently perform better with all thinning versions, all classes and for both discovery and validation cohorts. But there were cases, where the different feature selection and classifying methods seemed to give advantage when the data had been thinned. Such methods include the Fisher’s exact test, binary and multiclass ISPCA for DMR finding and feature selection and logistic regression model with DMR count variables.

## Acknowledgements

The calculations presented above were performed using computer resources within the Aalto University School of Science “Science-IT” project.

## Abbreviations

AML: acute myeloid leukemia
AUROC: area under receiver-operating characteristics curve
BLCA: bladder cancer
BRCA: breast cancer
BS-seq: bisulfite sequencing
cfDNA: cell-free DNA
cfMeDIP-seq: cell-free methylated DNA immunoprecipitation and high-throughput sequencing
CRC: colorectal cancer
ctDNA: circulating tumor DNA
DMC: differentially methylated cytosine
DMR: differentially methylated region
GO: gene ontology
GP: Gaussian process
ISPCA: iterative supervised principal component analysis
Lasso: least absolute shrinkage and selection operator
LR: logistic regression
LUC: lung cancer
MCMC: Markov chain Monte Carlo
NGS: next-generation sequencing
PCA: principal component analysis
PDAC: pancreatic ductal adenocarcinoma
RCC: renal cell carcinoma
RHS: regularized horseshoe
RRBS-seq: reduced representation bisulfite sequencing

## Availability of data and materials

Access to the cfMeDIP-seq data set was requested from the authors of [7]. Lists of the differentially methylated RRBS-seq based DMCs are available as Supplementary information of [7]. The scripts used to produce the results presented in this work are available at https://github.com/hallav/cfMeDIP-seq. Lists of found DMRs for moderated t-test, moderated t-test with new data transformation and Fisher’s exact test are available in Supplementary files 2, 3 and 4, respectively. The DMR files include the DMR location and number of data splits where the DMR in question was picked. The RPKM data for the intracranial tumors data set from [11] is available at the corresponding Zenodo archive [31].

## Authors’ contributions

VH and HL developed methods. VH performed the computational analysis and wrote the manuscript. HL participated in the revision of the manuscript. Both authors have read and accepted the manuscript.

## Footnotes

1 In case of overlapping DMRs, the dimension is less than 2100.

