## Supplementary file for "Probabilistic modeling methods for cell-free DNA methylation based cancer classification"

### Supplementary Information

Viivi Halla-aho and Harri Lähdesmäki

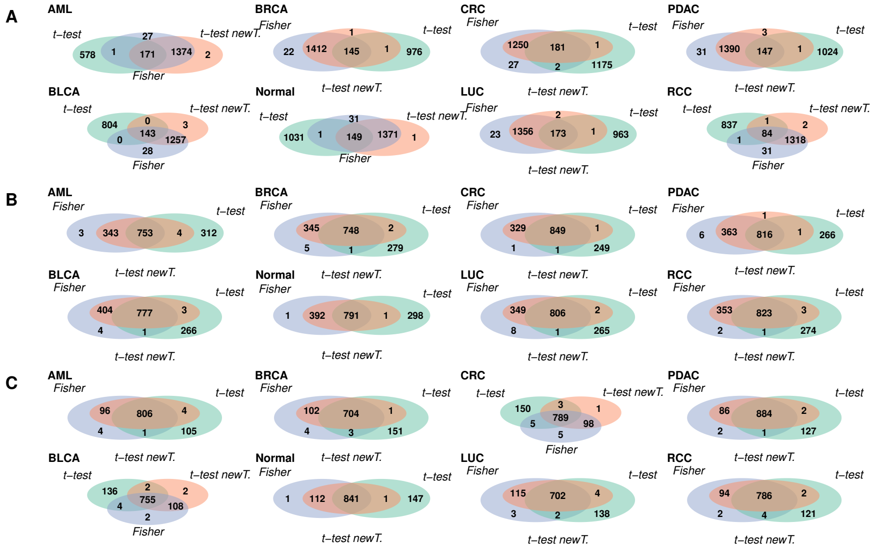


Figure S1: Number of overlapping DMRs between the DMRs found with Fisher's exact test, moderated t-test with the original data transformation and moderated t-test with new data transformation. DMRs from the 100 data splits were combined and only the DMRs present in the DMR set of at least 50 data splits were kept for finding the overlaps. The results are presented for thinning with total read counts $10^4$, $10^5$ and $10^6$ in figures **A**, **B** and **C**, respectively.

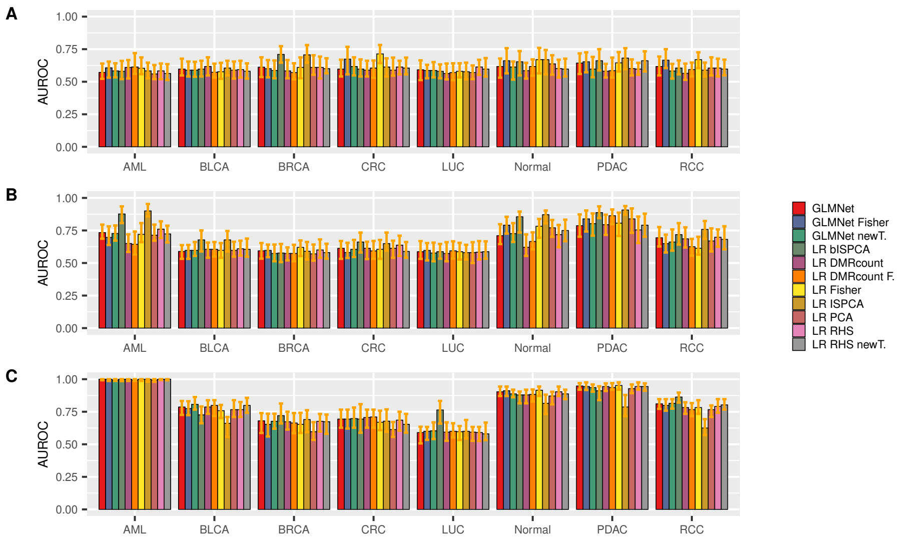


Figure S2: Barplots of the AUROC medians over the 100 data splits for all approaches. The orange bar on top of each bar shows the 25% and 75% quantiles. The AUROC values have been calculated for the test data sets in the discovery cohort. The results are presented for each class separately. The results are presented for thinning with total read counts $10^4$, $10^5$ and $10^6$ in figures **A**, **B** and **C**, respectively.

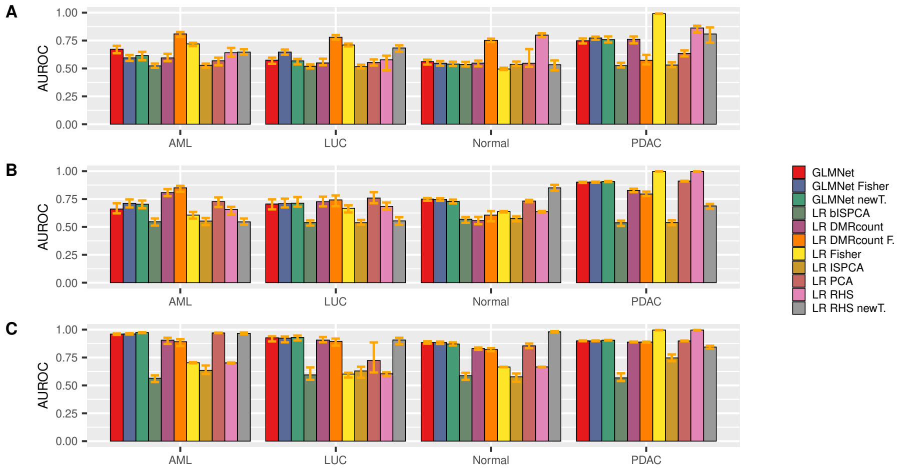


Figure S3: Barplot of the AUROC medians over the 100 data splits for all approaches on the validation cohort. The orange bar on top of each bar shows the 25% and 75% quantiles. The results are presented for each class separately. The results are presented for thinning with total read counts $10^4$, $10^5$ and $10^6$ in figures **A**, **B** and **C**, respectively.

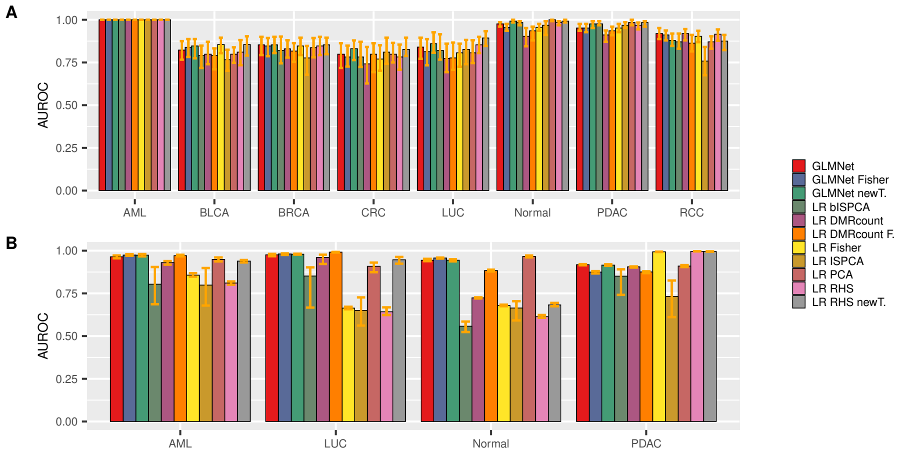


Figure S4: The bars represent the AUROC medians over the 100 data splits for the non-thinned data set for each of the methods. The orange bar on top of each bar shows the AUROC range from 25% quantile to 75% quantile. **A**: AUROC medians for discovery cohort. **B**: AUROC medians for validation cohort.

Table S1: Medians of AUROC and AUPRC values over 100 data splits for discovery cohort, subsampled data with total read count $10^4$. Highest values have been bolded.

| | | LR RHS | GLMNet | LR PCA | LR Fisher | LR DMRcount | LR DMRcount F. | LR lSPCA | LR blSPCA | LR RHS newT. | GLMNet newT. | GLMNet Fisher |
|---|---|---|---|---|---|---|---|---|---|---|---|---|
| AML | AUROC | 0.583 | 0.572 | 0.560 | 0.603 | 0.610 | **0.613** | 0.583 | 0.580 | 0.563 | 0.583 | 0.607 |
| | AUPRC | 0.149 | 0.150 | 0.138 | 0.130 | 0.120 | **0.263** | 0.159 | 0.155 | 0.138 | 0.158 | 0.160 |
| BLCA | AUROC | 0.593 | 0.597 | 0.589 | 0.577 | **0.617** | 0.573 | 0.605 | 0.597 | 0.581 | 0.589 | 0.589 |
| | AUPRC | **0.157** | 0.145 | 0.108 | 0.120 | 0.120 | 0.122 | 0.116 | 0.121 | 0.105 | 0.094 | 0.142 |
| BRCA | AUROC | 0.610 | 0.612 | 0.610 | 0.610 | 0.583 | 0.570 | 0.707 | **0.710** | 0.600 | 0.587 | 0.600 |
| | AUPRC | 0.197 | 0.188 | 0.142 | 0.188 | 0.159 | 0.158 | 0.271 | **0.280** | 0.160 | 0.159 | 0.190 |
| CRC | AUROC | 0.613 | 0.597 | 0.589 | **0.714** | 0.589 | 0.605 | 0.613 | 0.597 | 0.605 | 0.617 | 0.673 |
| | AUPRC | 0.150 | 0.127 | 0.103 | 0.247 | 0.111 | 0.128 | 0.150 | 0.154 | 0.096 | 0.093 | **0.279** |
| LUC | AUROC | **0.613** | 0.592 | 0.570 | 0.580 | 0.563 | 0.570 | 0.577 | 0.580 | 0.597 | 0.587 | 0.587 |
| | AUPRC | 0.120 | 0.134 | 0.132 | 0.160 | 0.141 | 0.145 | **0.192** | 0.172 | 0.128 | 0.142 | 0.153 |
| Normal | AUROC | 0.601 | 0.617 | 0.637 | **0.669** | 0.585 | 0.617 | **0.669** | 0.653 | 0.597 | 0.613 | 0.661 |
| | AUPRC | 0.107 | 0.113 | 0.098 | **0.237** | 0.109 | 0.120 | 0.199 | 0.185 | 0.119 | 0.111 | 0.225 |
| PDAC | AUROC | 0.597 | 0.643 | 0.601 | 0.645 | 0.581 | 0.585 | **0.681** | 0.661 | 0.661 | 0.597 | 0.653 |
| | AUPRC | 0.129 | 0.158 | 0.110 | **0.243** | 0.125 | 0.105 | 0.239 | 0.235 | 0.191 | 0.139 | 0.227 |
| RCC | AUROC | 0.605 | 0.617 | 0.605 | **0.669** | 0.565 | 0.593 | 0.589 | 0.605 | 0.597 | 0.583 | 0.665 |
| | AUPRC | 0.102 | 0.099 | 0.122 | **0.185** | 0.108 | 0.114 | 0.123 | 0.124 | 0.107 | 0.113 | **0.185** |

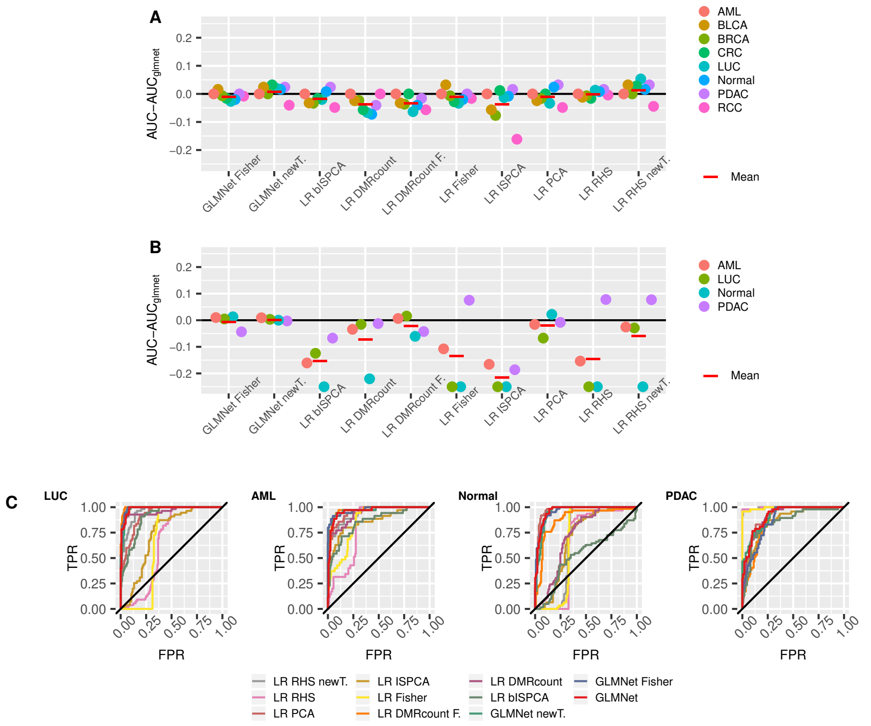


Figure S5: Assessing the performance of the different classifiers for the non-thinned data. **A**: The median AUROC differences between the original GLMnet method and the other classifiers for the discovery cohort test sets. The median has been calculated over the 100 data splits. **B**: The median AUROC differences between the original GLMnet method and the other classifiers for the validation cohort data. The median has been calculated over the 100 data splits. The negative values have been truncated to $-0.25$. **C**: Validation cohort ROCs calculated with prediction means over 100 data splits and corresponding fitted models. The results are presented for each class separately. Red lines indicate the means over the four classes.

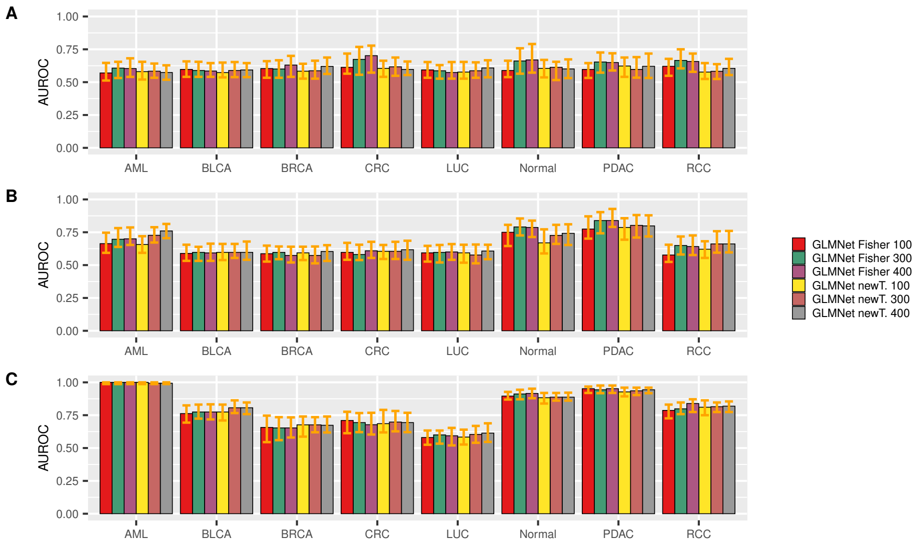


Figure S6: Comparison of different DMR numbers. AUROC barplots for the discovery cohort. The orange bar on top of each bar shows the 25% and 75% quantiles. The results are presented for thinning with total read counts $10^4$, $10^5$ and $10^6$ in figures **A**, **B** and **C**, respectively.

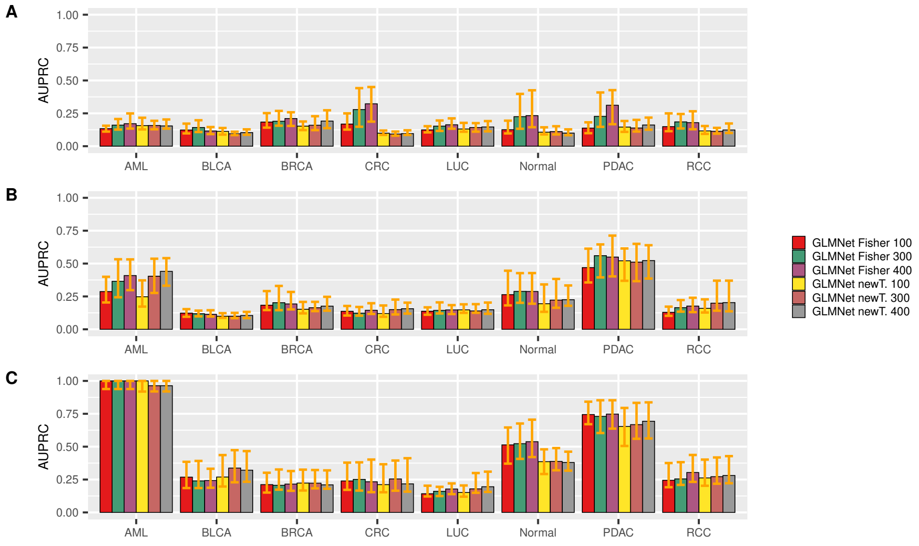


Figure S7: Comparison of different DMR numbers. AUPRC barplot for the discovery cohort. The orange bar on top of each bar shows the 25% and 75% quantiles. The results are presented for thinning with total read counts $10^4$, $10^5$ and $10^6$ in figures **A**, **B** and **C**, respectively.

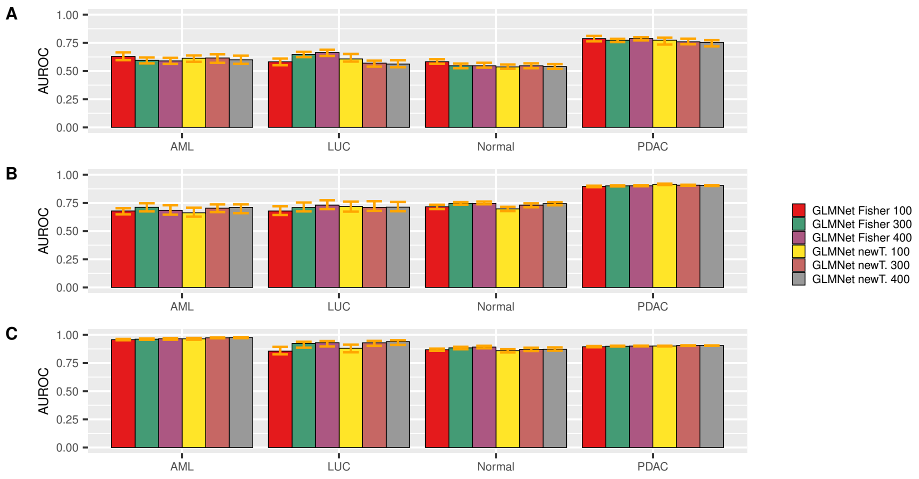


Figure S8: Comparison of different DMR numbers. AUROC barplot for the validation cohort. The orange bar on top of each bar shows the 25% and 75% quantiles. The results are presented for thinning with total read counts $10^4$, $10^5$ and $10^6$ in figures **A**, **B** and **C**, respectively.

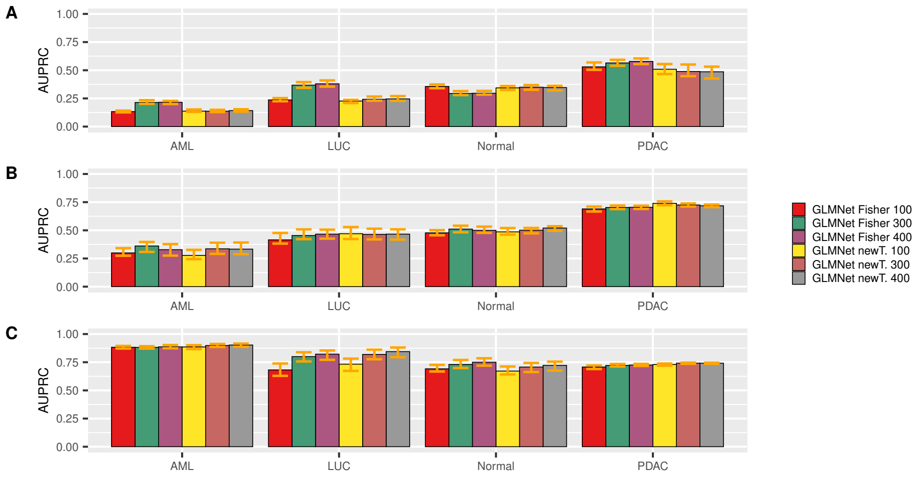


Figure S9: Comparison of different DMR numbers. AUPRC barplot for the validation cohort. The orange bar on top of each bar shows the 25% and 75% quantiles. The results are presented for thinning with total read counts $10^4$, $10^5$ and $10^6$ in figures **A**, **B** and **C**, respectively.

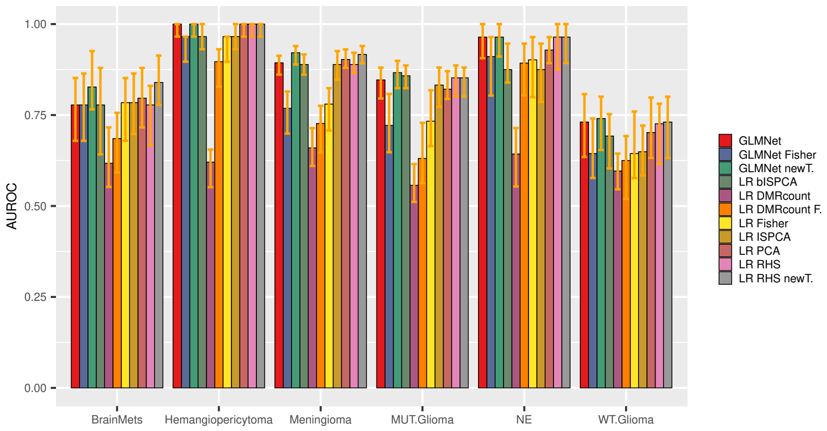


Figure S10: he bars represent the AUROC medians over the 100 data splits for the intracranial tumors data set for each of the methods. The orange bar on top of each bar shows the AUROC range from 25% quantile to 75% quantile. Abbreviations of the class names: BrainMets = brain metastases, MUT.Glioma= IDH mutant glioma, NE= low-grade glioneuronal and WT.Glioma= IDH wildtype glioma.

Table S2: Medians of AUROC and AUPRC values over 100 data splits for discovery cohort, subsampled data with total read count $10^5$. Highest values have been bolded.

| | | LR RHS | GLMNet | LR PCA | LR Fisher | LR DMRcount | LR DMRcount F. | LR lSPCA | LR bISPCA | LR RHS newT. | GLMNet newT. | GLMNet Fisher |
|---|---|---|---|---|---|---|---|---|---|---|---|---|
| AML | AUROC | 0.760 | 0.733 | 0.713 | 0.720 | 0.650 | 0.643 | **0.900** | 0.877 | 0.723 | 0.727 | 0.697 |
| | AUPRC | 0.470 | 0.422 | 0.394 | 0.412 | 0.233 | 0.222 | **0.703** | 0.699 | 0.379 | 0.404 | 0.365 |
| BLCA | AUROC | 0.609 | 0.589 | 0.597 | 0.597 | 0.605 | 0.605 | **0.677** | **0.677** | 0.605 | 0.597 | 0.597 |
| | AUPRC | 0.092 | 0.104 | 0.102 | 0.113 | 0.113 | 0.127 | **0.191** | 0.174 | 0.106 | 0.100 | 0.119 |
| BRCA | AUROC | 0.600 | 0.593 | 0.570 | **0.620** | 0.573 | 0.573 | 0.590 | 0.577 | 0.580 | 0.573 | 0.597 |
| | AUPRC | 0.211 | 0.198 | 0.166 | 0.220 | 0.158 | 0.144 | 0.175 | 0.176 | 0.168 | 0.164 | 0.202 |
| CRC | AUROC | 0.637 | 0.613 | 0.613 | 0.605 | 0.613 | 0.593 | 0.649 | **0.661** | 0.589 | 0.605 | 0.581 |
| | AUPRC | 0.161 | 0.145 | 0.152 | 0.133 | 0.126 | 0.121 | **0.284** | 0.260 | 0.137 | 0.150 | 0.123 |
| LUC | AUROC | 0.583 | 0.587 | 0.580 | 0.587 | 0.573 | 0.593 | 0.580 | 0.587 | 0.587 | 0.577 | **0.597** |
| | AUPRC | 0.141 | 0.147 | 0.139 | 0.146 | 0.152 | **0.160** | 0.157 | 0.151 | 0.129 | 0.140 | 0.144 |
| Normal | AUROC | 0.718 | 0.710 | 0.770 | 0.782 | 0.621 | 0.665 | **0.871** | 0.855 | 0.75 | 0.726 | 0.79 |
| | AUPRC | 0.210 | 0.219 | 0.250 | 0.318 | 0.163 | 0.177 | **0.337** | 0.307 | 0.256 | 0.222 | 0.288 |
| PDAC | AUROC | 0.754 | 0.786 | 0.839 | 0.806 | 0.794 | 0.863 | **0.907** | 0.887 | 0.79 | 0.802 | 0.839 |
| | AUPRC | 0.446 | 0.490 | 0.566 | 0.502 | 0.540 | 0.610 | **0.643** | 0.630 | 0.531 | 0.511 | 0.560 |
| RCC | AUROC | 0.698 | 0.694 | 0.669 | 0.613 | 0.685 | 0.625 | **0.758** | 0.718 | 0.681 | 0.661 | 0.649 |
| | AUPRC | 0.206 | 0.201 | 0.210 | 0.156 | 0.215 | 0.152 | **0.238** | 0.230 | 0.234 | 0.199 | 0.165 |

Table S3: Medians of AUROC and AUPRC values over 100 data splits for discovery cohort, subsampled data with total read count $10^6$. Highest values have been bolded.

| | | LR RHS | GLMNet | LR PCA | LR Fisher | LR DMRcount | LR DMRcount F. | LR ISPCA | LR bISPCA | LR RHS newT. | GLMNet newT. | GLMNet Fisher |
|---|---|---|---|---|---|---|---|---|---|---|---|---|
| AML | AUROC | **1.000** | **1.000** | 0.993 | **1.000** | **1.000** | 0.993 | **1.000** | **1.000** | **1.000** | 0.993 | **1.000** |
| | AUPRC | **1.000** | **1.000** | 0.964 | **1.000** | **1.000** | 0.964 | **1.000** | **1.000** | **1.000** | 0.964 | **1.000** |
| BLCA | AUROC | 0.766 | 0.786 | 0.766 | 0.758 | 0.786 | 0.798 | 0.661 | 0.726 | 0.798 | **0.806** | 0.774 |
| | AUPRC | 0.253 | 0.253 | 0.237 | 0.235 | 0.260 | 0.262 | 0.182 | 0.209 | **0.338** | 0.337 | 0.240 |
| BRCA | AUROC | 0.677 | 0.680 | 0.597 | 0.653 | 0.673 | 0.663 | 0.690 | **0.720** | 0.673 | 0.677 | 0.653 |
| | AUPRC | 0.233 | 0.224 | 0.168 | 0.207 | 0.227 | 0.231 | **0.309** | 0.322 | 0.239 | 0.222 | 0.206 |
| CRC | AUROC | 0.685 | 0.694 | 0.613 | 0.669 | 0.706 | **0.710** | 0.677 | 0.694 | 0.653 | 0.698 | 0.694 |
| | AUPRC | 0.202 | 0.223 | 0.143 | 0.225 | 0.231 | 0.242 | 0.196 | 0.383 | 0.173 | **0.254** | 0.251 |
| LUC | AUROC | 0.590 | 0.590 | 0.593 | 0.597 | 0.593 | 0.600 | 0.600 | **0.763** | 0.580 | 0.603 | 0.600 |
| | AUPRC | 0.150 | 0.157 | 0.165 | 0.155 | 0.171 | 0.166 | 0.185 | **0.431** | 0.160 | 0.179 | 0.160 |
| Normal | AUROC | 0.903 | 0.903 | 0.871 | **0.915** | 0.879 | 0.883 | 0.815 | 0.879 | 0.887 | 0.887 | 0.911 |
| | AUPRC | 0.533 | 0.522 | 0.375 | **0.544** | 0.430 | 0.474 | 0.360 | 0.408 | 0.372 | 0.388 | 0.523 |
| PDAC | AUROC | 0.944 | **0.948** | 0.927 | 0.952 | 0.944 | 0.935 | 0.786 | 0.911 | 0.944 | 0.935 | 0.944 |
| | AUPRC | 0.736 | 0.736 | 0.660 | **0.762** | 0.733 | 0.723 | 0.425 | 0.667 | 0.673 | 0.667 | 0.730 |
| RCC | AUROC | 0.790 | 0.810 | 0.766 | 0.782 | 0.782 | 0.766 | 0.625 | **0.863** | 0.802 | 0.815 | 0.798 |
| | AUPRC | 0.277 | 0.281 | 0.204 | 0.249 | 0.242 | 0.214 | 0.147 | **0.319** | 0.266 | 0.272 | 0.255 |

Table S4: Medians of AUROC and AUPRC values over 100 data splits for validation cohort, subsampled data with total read count $10^4$. Highest values have been bolded.

| | | LR RHS | GLMNet | LR PCA | LR Fisher | LR DMRcount | LR DMRcount F. | LR ISPCA | LR bISPCA | GLMNet newT. | GLMNet Fisher | LR RHS newT. |
|---|---|---|---|---|---|---|---|---|---|---|---|---|
| AML | AUROC | 0.641 | 0.672 | 0.569 | 0.720 | 0.595 | **0.809** | 0.528 | 0.523 | 0.595 | 0.616 | 0.646 |
| | AUPRC | 0.131 | 0.126 | 0.148 | 0.116 | 0.137 | **0.411** | 0.198 | 0.200 | 0.214 | 0.136 | 0.133 |
| LUC | AUROC | 0.578 | 0.574 | 0.553 | 0.710 | 0.550 | **0.780** | 0.516 | 0.520 | 0.647 | 0.570 | 0.683 |
| | AUPRC | 0.266 | 0.244 | 0.250 | 0.190 | 0.254 | **0.489** | 0.282 | 0.281 | 0.368 | 0.243 | 0.223 |
| Normal | AUROC | **0.799** | 0.561 | 0.546 | 0.495 | 0.548 | 0.752 | 0.537 | 0.536 | 0.547 | 0.546 | 0.534 |
| | AUPRC | **0.598** | 0.374 | 0.374 | 0.280 | 0.342 | 0.207 | 0.294 | 0.299 | 0.294 | 0.349 | 0.349 |
| PDAC | AUROC | 0.863 | 0.749 | 0.635 | **0.991** | 0.761 | 0.573 | 0.531 | 0.525 | 0.773 | 0.759 | 0.809 |
| | AUPRC | 0.144 | 0.453 | 0.319 | **0.984** | 0.529 | 0.211 | 0.238 | 0.237 | 0.565 | 0.489 | 0.406 |

Table S5: Medians of AUROC and AUPRC values over 100 data splits for validation cohort, subsampled data with total read count $10^5$. Highest values have been bolded.

| | | LR RHS | GLMNet | LR PCA | LR Fisher | LR DMRcount | LR DMRcount F. | LR ISPCA | LR bISPCA | GLMNet newT. | GLMNet Fisher | LR RHS newT. |
|---|---|---|---|---|---|---|---|---|---|---|---|---|
| AML | AUROC | 0.657 | 0.660 | 0.727 | 0.606 | 0.808 | **0.850** | 0.552 | 0.547 | 0.711 | 0.703 | 0.546 |
| | AUPRC | 0.125 | 0.296 | 0.360 | 0.134 | 0.482 | **0.552** | 0.210 | 0.195 | 0.361 | 0.336 | 0.165 |
| LUC | AUROC | 0.684 | 0.705 | **0.759** | 0.666 | 0.726 | 0.742 | 0.538 | 0.539 | 0.709 | 0.711 | 0.554 |
| | AUPRC | 0.195 | 0.452 | **0.516** | 0.199 | 0.448 | 0.445 | 0.312 | 0.309 | 0.454 | 0.464 | 0.297 |
| Normal | AUROC | 0.637 | 0.749 | 0.732 | 0.638 | 0.557 | 0.606 | 0.577 | 0.569 | 0.747 | 0.73 | **0.850** |
| | AUPRC | 0.349 | 0.511 | 0.456 | 0.350 | 0.277 | 0.254 | 0.371 | 0.362 | 0.510 | 0.501 | **0.811** |
| PDAC | AUROC | **0.998** | 0.901 | 0.911 | **0.998** | 0.827 | 0.795 | 0.539 | 0.538 | 0.902 | 0.907 | 0.688 |
| | AUPRC | 0.993 | 0.699 | 0.739 | **0.994** | 0.660 | 0.652 | 0.247 | 0.243 | 0.703 | 0.725 | 0.297 |

Table S6: Medians of AUROC and AUPRC values over 100 data splits for validation cohort, subsampled data with total read count $10^6$. Highest values have been bolded.

| | | LR RHS | GLMNet | LR PCA | LR Fisher | LR DMRcount | LR DMRcount F. | LR ISPCA | LR bISPCA | GLMNet newT. | GLMNet Fisher | LR RHS newT. |
|---|---|---|---|---|---|---|---|---|---|---|---|---|
| AML | AUROC | 0.702 | 0.960 | 0.970 | 0.703 | 0.905 | 0.892 | 0.635 | 0.564 | 0.961 | **0.974** | 0.966 |
| | AUPRC | 0.245 | 0.883 | 0.875 | 0.245 | 0.798 | 0.747 | 0.233 | 0.198 | 0.882 | 0.898 | **0.906** |
| LUC | AUROC | 0.604 | 0.927 | 0.723 | 0.601 | 0.906 | 0.895 | 0.629 | 0.594 | 0.923 | **0.929** | 0.907 |
| | AUPRC | 0.295 | 0.815 | 0.271 | 0.294 | 0.781 | 0.742 | 0.370 | 0.359 | 0.800 | **0.819** | 0.781 |
| Normal | AUROC | 0.666 | 0.884 | 0.855 | 0.665 | 0.830 | 0.821 | 0.577 | 0.587 | 0.884 | 0.870 | **0.981** |
| | AUPRC | 0.367 | 0.739 | 0.675 | 0.367 | 0.626 | 0.605 | 0.267 | 0.259 | 0.730 | 0.707 | **0.954** |
| PDAC | AUROC | **0.996** | 0.900 | 0.899 | **0.996** | 0.888 | 0.888 | 0.747 | 0.568 | 0.899 | 0.905 | 0.844 |
| | AUPRC | **0.992** | 0.724 | 0.712 | 0.991 | 0.706 | 0.706 | 0.525 | 0.210 | 0.721 | 0.742 | 0.538 |

Table S7: Medians of AUROC and AUPRC values over 100 data splits for discovery cohort, non-thinned data. Highest values have been bolded.

| | | LR RHS | GLMNet | LR PCA | LR Fisher | LR DMRcount | LR DMRcount F. | LR ISPCA | LR bISPCA | GLMNet newT. | GLMNet Fisher | LR RHS newT. |
|---|---|---|---|---|---|---|---|---|---|---|---|---|
| AML | AUROC | **1.000** | **1.000** | **1.000** | **1.000** | **1.000** | **1.000** | **1.000** | **1.000** | **1.000** | **1.000** | **1.000** |
| | AUPRC | **1.000** | **1.000** | **1.000** | **1.000** | **1.000** | **1.000** | **1.000** | **1.000** | **1.000** | **1.000** | **1.000** |
| BLCA | AUROC | 0.810 | 0.823 | 0.798 | **0.855** | 0.798 | 0.790 | 0.766 | 0.790 | 0.839 | 0.847 | **0.855** |
| | AUPRC | 0.360 | 0.380 | 0.282 | 0.396 | 0.321 | 0.225 | 0.350 | 0.265 | 0.323 | 0.435 | **0.447** |
| BRCA | AUROC | 0.847 | **0.853** | 0.837 | 0.847 | 0.830 | 0.817 | 0.777 | 0.820 | 0.847 | **0.853** | **0.853** |
| | AUPRC | **0.558** | 0.521 | 0.484 | 0.534 | 0.493 | 0.444 | 0.529 | 0.549 | 0.518 | 0.532 | 0.547 |
| CRC | AUROC | 0.782 | 0.798 | 0.798 | 0.770 | 0.742 | 0.798 | 0.810 | 0.782 | 0.782 | **0.831** | 0.827 |
| | AUPRC | 0.582 | 0.580 | 0.560 | 0.585 | 0.563 | 0.568 | 0.580 | 0.576 | 0.583 | 0.587 | **0.599** |
| LUC | AUROC | 0.853 | 0.840 | 0.807 | 0.807 | 0.773 | 0.777 | 0.827 | 0.820 | 0.813 | 0.860 | **0.893** |
| | AUPRC | 0.594 | 0.662 | 0.589 | 0.507 | 0.433 | 0.368 | 0.570 | 0.665 | 0.532 | **0.743** | 0.735 |
| Normal | AUROC | 0.984 | 0.976 | **1.000** | 0.956 | 0.903 | 0.935 | 0.968 | 0.984 | 0.956 | 0.992 | 0.992 |
| | AUPRC | 0.909 | 0.837 | **1.000** | 0.744 | 0.561 | 0.562 | 0.797 | 0.872 | 0.736 | 0.944 | 0.944 |
| PDAC | AUROC | 0.968 | 0.952 | **0.984** | 0.952 | 0.911 | 0.935 | 0.968 | 0.976 | 0.952 | 0.976 | **0.984** |
| | AUPRC | 0.842 | 0.748 | **0.909** | 0.758 | 0.541 | 0.699 | 0.812 | 0.872 | 0.751 | 0.853 | 0.884 |
| RCC | AUROC | 0.915 | 0.919 | 0.871 | 0.903 | **0.919** | 0.863 | 0.758 | 0.871 | 0.911 | 0.879 | 0.875 |
| | AUPRC | 0.477 | 0.535 | 0.350 | 0.477 | **0.559** | 0.400 | 0.239 | 0.373 | 0.518 | 0.398 | 0.335 |

Table S8: Medians of AUROC and AUPRC values over 100 data splits for validation cohort, non-thinned data. Highest values have been bolded.

| | | LR RHS | GLMNet | LR PCA | LR Fisher | LR DMRcount | LR DMRcount F. | LR ISPCA | LR bISPCA | GLMNet newT. | GLMNet Fisher | LR RHS newT. |
|---|---|---|---|---|---|---|---|---|---|---|---|---|
| AML | AUROC | 0.811 | 0.964 | 0.948 | 0.857 | 0.930 | 0.971 | 0.799 | 0.804 | **0.974** | **0.974** | 0.939 |
| | AUPRC | 0.402 | 0.807 | 0.865 | 0.581 | 0.791 | 0.902 | 0.577 | 0.554 | **0.938** | 0.933 | 0.843 |
| LUC | AUROC | 0.642 | 0.976 | 0.909 | 0.664 | 0.960 | **0.991** | 0.650 | 0.852 | 0.981 | 0.979 | 0.947 |
| | AUPRC | 0.316 | 0.932 | 0.784 | 0.329 | 0.901 | **0.976** | 0.219 | 0.669 | 0.945 | 0.940 | 0.869 |
| Normal | AUROC | 0.614 | 0.944 | **0.966** | 0.679 | 0.724 | 0.884 | 0.664 | 0.558 | 0.957 | 0.944 | 0.683 |
| | AUPRC | 0.342 | 0.853 | **0.911** | 0.372 | 0.453 | 0.712 | 0.247 | 0.351 | 0.900 | 0.847 | 0.374 |
| PDAC | AUROC | **0.996** | 0.918 | 0.910 | 0.993 | 0.906 | 0.875 | 0.732 | 0.851 | 0.875 | 0.915 | 0.995 |
| | AUPRC | **0.992** | 0.783 | 0.760 | 0.986 | 0.664 | 0.627 | 0.478 | 0.626 | 0.669 | 0.803 | 0.989 |

Table S9: Medians of the AUROC and AUPRC values over 100 data splits for intracranial tumors data set. Highest values have been bolded.

| | | LR RHS | GLMNet | LR PCA | LR Fisher | LR DMRcount | LR DMRcount F. | LR ISPCA | LR bISPCA | GLMNet Fisher | GLMNet newT. | LR RHS newT. |
|---|---|---|---|---|---|---|---|---|---|---|---|---|
| Brain metastases | AUROC | 0.778 | 0.778 | 0.796 | 0.784 | 0.617 | 0.685 | 0.784 | 0.778 | 0.778 | 0.827 | **0.840** |
| | AUPRC | 0.226 | 0.311 | 0.273 | 0.324 | 0.131 | 0.186 | 0.337 | 0.302 | 0.348 | **0.488** | 0.480 |
| Hemangiopericytoma | AUROC | **1.000** | **1.000** | **1.000** | 0.966 | 0.621 | 0.897 | 0.966 | 0.966 | 0.966 | **1.000** | **1.000** |
| | AUPRC | **1.000** | **1.000** | **1.000** | 0.307 | 0.038 | 0.137 | 0.307 | 0.307 | 0.307 | **1.000** | **1.000** |
| Meningioma | AUROC | 0.889 | 0.894 | 0.903 | 0.780 | 0.660 | 0.727 | 0.889 | 0.889 | 0.769 | **0.921** | 0.917 |
| | AUPRC | 0.844 | 0.859 | 0.852 | 0.746 | 0.665 | 0.690 | 0.813 | 0.830 | 0.724 | **0.874** | 0.869 |
| Low-grade glioneuronal | AUROC | **0.964** | **0.964** | 0.929 | 0.902 | 0.643 | 0.893 | 0.875 | 0.875 | 0.911 | **0.964** | **0.964** |
| | AUPRC | 0.689 | **0.712** | 0.615 | 0.451 | 0.085 | 0.491 | 0.234 | 0.451 | 0.544 | **0.712** | **0.712** |
| IDH wildtype glioma | AUROC | 0.726 | 0.731 | 0.702 | 0.644 | 0.596 | 0.625 | 0.649 | 0.692 | 0.644 | **0.740** | 0.731 |
| | AUPRC | 0.262 | 0.277 | 0.275 | **0.283** | 0.137 | 0.153 | 0.216 | 0.203 | 0.240 | 0.257 | 0.257 |
| IDH mutant glioma | AUROC | 0.852 | 0.847 | 0.821 | 0.733 | 0.557 | 0.631 | 0.832 | 0.858 | 0.722 | **0.866** | 0.852 |
| | AUPRC | 0.626 | 0.615 | 0.597 | 0.524 | 0.268 | 0.418 | 0.588 | 0.595 | 0.502 | **0.637** | 0.627 |
